# Dynamic estimation of metabolic state during CAR T cell production and the relationship of early metabolism to final therapeutic product

**DOI:** 10.1101/2025.05.11.653250

**Authors:** N Suhas Jagannathan, Wei-Xiang Sin, Denise Bei Lin Teo, Faris Kairi, Yen Hoon Luah, Francesca Lorraine Wei Inng Lim, Michaela Su-Fern Seng, Shui Yen Soh, Yie Hou Lee, Lisa Tucker-Kellogg, Michael E. Birnbaum, Rajeev J. Ram

## Abstract

Adoptive cell therapies such as CAR T cells have revolutionized cancer treatment and shown successes even with refractory cases of haematological malignancies. There is burgeoning interest in the optimization and improved manufacturing of cell therapy products. For CAR T cell therapy, enrichment of certain phenotypes of T cells in the infusion product have been correlated with improved long-term treatment outcomes. While metabolic control of T cell phenotypic fates has been demonstrated in some contexts, gaps still exist in our knowledge of how T cell metabolic dynamics early during CAR T manufacturing might affect critical quality attributes (CQAs) of the final product (e.g., differentiation/exhaustion/potency). We present a modelling framework that can perform real-time estimation of per-cell metabolic rates of T cells expanded *ex vivo* in a reactor. We validate our estimated rates using metabolic assays, show how average rates can be deconvoluted to rates of individual T cell phenotypes, and demonstrate applicability to different reactor types. Applying our tool to the expansion of both healthy and patient-derived cells in a perfusion-based microbioreactor, we offer proof-of-principle to show that correlations exist between early metabolic rates of T cells in culture and cellular attributes related to growth, differentiation and exhaustion of the final product. Given the biological variation that exists in the growth and dynamics of patient-derived cells in culture, such modelling contributes to the overarching goal of improving the consistency of cell therapy through Adaptive Process Control (APC).

## Introduction

Adoptive cell therapies have provided a transformative approach to treatments in the spheres of cancer and regenerative medicine, sometimes as the last resort where other therapeutic modalities have faced limited success^1^. Specifically for cancer immunotherapy, paradigms such as CAR T cell therapy (seven FDA-approved products for haematological cancers^2,3^), T cell receptor therapy (one FDA-approved product for sarcoma^4^) and CAR NK-therapy (in clinical trials^5^) are revolutionizing treatment by leveraging the host’s immune system to target and eliminate malignant cells, with successful remission for over 10 years^6,7^. Consequently, there is increasing interest in the manufacturing and optimization of autologous cell therapy products^8–11^. However, unlike conventional biologics, the patient-specific nature of autologous therapies mandates a customized/adaptive manufacturing protocol rather than a one-size-fits-all approach. This is a consequence of high inter-patient variabilities in treatment history, patient demography and disease progression that can affect cellular states (proliferation/differentiation/metabolism) of both patient-derived source cells and the final cell therapy product.

For current autologous CAR T cell therapies, conventional manufacturing involves PBMC extraction from patients, purification and activation of T cells, viral transduction of the CAR construct, and finally cellular expansion to clinical dosage levels of CAR T cells in a bioreactor. The efficacy of CAR T cells depends both on characteristics of the source T cells and on the phenotypes of the expanded T cells^12^. Early identification of the phenotypic/metabolic state of cells while in *ex vivo* culture can aid in-process monitoring and control of population dynamics, greatly augmenting our ability to tailor CAR T therapy to each patient (Figure 1).

**Figure 1.**
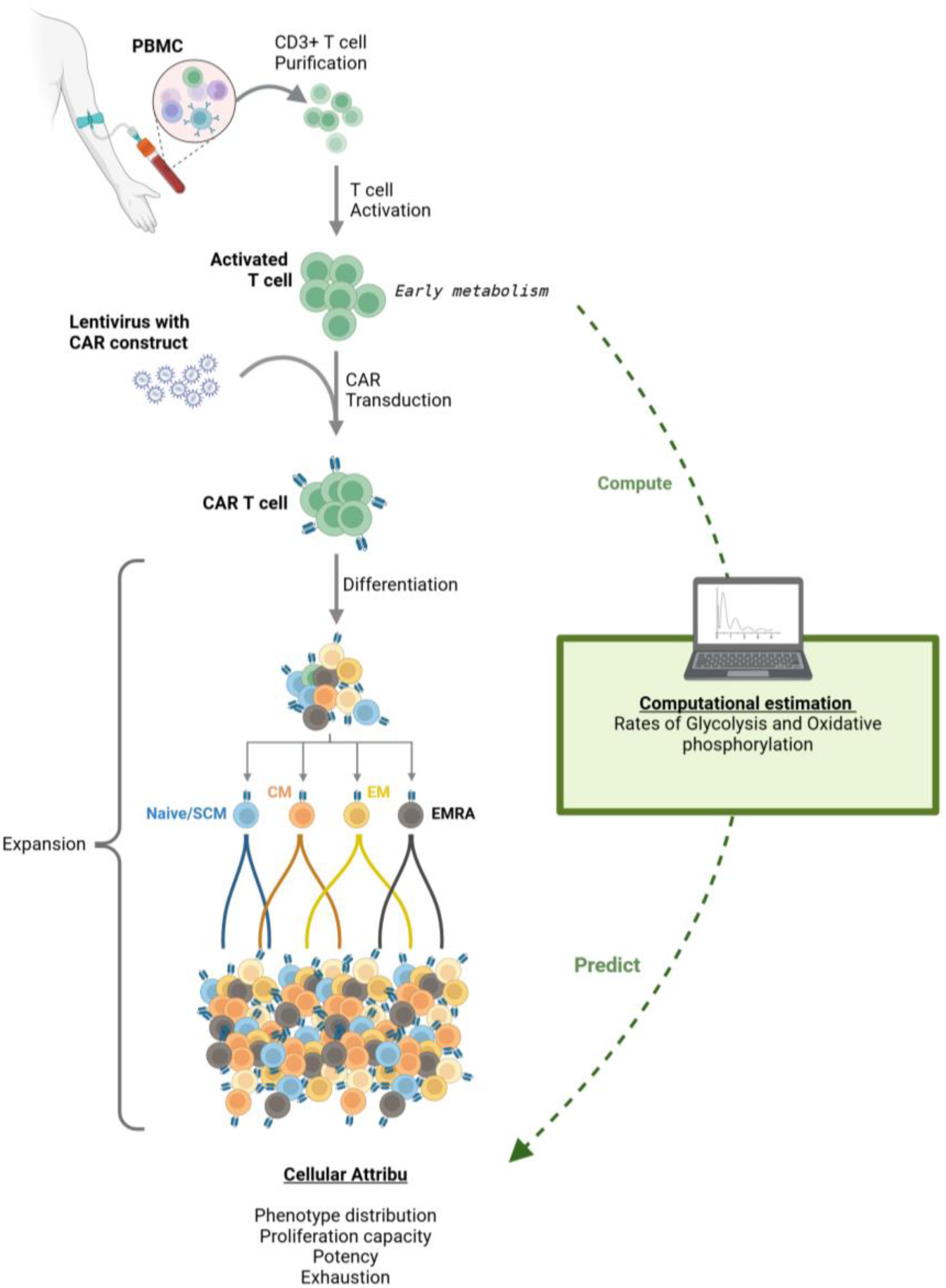
CAR T cell growth and dynamics during the manufacturing process. PBMCs extracted from patients are further purified to extract T cells which are activated, transduced with the CAR construct and expanded in a reactor to target cell densities. Activation, T cell differentiation and exhaustion all alter T cell metabolism, especially the rates of glycolysis and oxidative phosphorylation (OxPhos). Understanding relationships between early metabolic trends and final critical quality attributes (CQAs) such as potency, phenotypic distribution and exhaustion is the first step towards metabolic control of the CAR T cell manufacturing process. Created in BioRender. Jagannathan, S. (2025) https://BioRender.com/e44d2ca

Tracking phenotypic dynamics is conventionally performed through offline experimental characterization of cells sampled from the bioreactor. However, frequent sampling of cells is often infeasible, and poses sterility concerns for clinical GMP manufacturing. Metabolism can thus be an indirect monitor of phenotype dynamics, as it is well known that most phenotypic changes that occur during T cell activation or differentiation are accompanied by quantifiable changes in metabolic state^13^. Of note, relative activity of oxidative phosphorylation (OxPhos) and aerobic glycolysis (glucose uptake followed by lactate export) can be instructive of cellular phenotypic states. In the physiological setting, T cells that have not encountered antigen are typically of the naïve phenotype (T_N_) –quiescent cells that depend primarily on a basal level of OxPhos for maintenance and self-renewal^14,15^. Upon antigen stimulation or activation, naïve cells differentiate into effector cells (T_EFF_) that are highly proliferative and show increased OxPhos and aerobic glycolysis^14,16–19^. Although T_EFF_ cells display high killing potency their short-lived nature makes them unsuitable for durable therapeutic purposes. Following activation, a subset of T cells differentiate into long-lived memory phenotypes that exhibit different metabolic profiles based on their subtype^13,20–22^. Central memory (T_CM_) and Stem cell memory (T_SCM_) cells are more OxPhos dependent than T_EFF_ cells. Effector memory cells (T_EM_) show higher levels of aerobic glycolysis than other memory cells but less than T_EFF_ cells ^19^. Terminally differentiated memory cells (T_EMRA_) are senescent cells that display T_EFF_ like potency^23,24^. CD4+ T_EMRA_ cells show greater dependence on OxPhos while CD8+ T_EMRA_ cells show increased aerobic glycolysis^25^. While T_EFF_, T_EM_ and T_EMRA_ cells are the most potent, T_N_, T_SCM_ and T_CM_ are preferred for infusion as they show increased survival in the tumour microenvironment and provide long lasting therapeutic benefits^26–29^.

Metabolic control of T cell fate has been demonstrated by other studies, through molecular targeting of metabolic enzymes or genetic manipulation. For example, inducing glycolysis during T cell differentiation can bias the population towards T_EM_^30^, while suppressing glycolysis during activation results in more but slower proliferating memory subtypes^31^. Beyond affecting differentiation, studies have shown that altering metabolism during expansion can affect the potency and persistence of *ex vivo* expanded CAR T cells. For example, increasing glucose uptake through overexpression of the GLUT1 transporter was found to increase CAR T potency^32^, while increasing mitochondrial activity was found to improve CAR T persistence^33^. These observations lend credence to the hypothesis that early identification of T cell metabolic states in the reactor during expansion can help direct manufacturing towards desirable phenotypic outcomes via metabolic control.

Identification of cellular states (State estimation) is an established area of research in computational modelling. Metabolic state estimation using sparse metabolite measurements and advanced techniques such as Kalman filters, neural networks, statistical models and metabolic flux analysis has been previously employed for bacterial/yeast cultures, antibody production etc.^34–36^. However most such cases involving simpler organisms do not face problems that apply to mammalian cell cultures, such as dynamically changing phenotypic/differentiation states of cells. Thus, there exists a need for a computational tool that can estimate metabolic and phenotypic states of dynamically changing systems such as CAR T cells during expansion in a bioreactor. Here, we develop a framework that integrates sparse metabolite measurements with metabolic modelling to estimate the metabolic state of T cells, paving the way for adaptive process control (APC). Many production processes have been radically improved using adaptive process control. APC is organized around analysis of non-destructive monitoring of the production process, especially continuous real-time monitoring. However, relatively few characteristics of cell culture are amenable to continuous real-time monitoring (e.g., pH, gas fraction), and the correlation between these available measurements and the product quality is unknown. If non-destructive measurements can used for estimating the cell phenotype (e.g., fate, quality), then subsequent studies can use existing knowledge of cell fate influences to adjust process variables (e.g., alter the culture media in real-time) for the purpose of improving product quality, as evidenced by the real-time monitoring. APC is a long-term goal with powerful promise for improvements of any bioprocess, but the key technology necessary for unlocking further APC research is the ability to correlate available non-destructive measurements with meaningful attributes of cell phenotype, fate, or quality.

In this work, we develop a computational model that uses real-time on-line process monitoring readouts and off-line metabolite measurements of perfusate samples as inputs to computationally estimate the cellular metabolic states in the culture over time. Our modelling method is particularly focused on glucose and lactate metabolism, as these indicate the relative rates of glycolysis and OxPhos, central pathways that can affect T cell differentiation. We previously demonstrated proof-of-principle capabilities of our tool in estimating rates of growth, oxygen consumption (as surrogate measurement to OxPhos) and lactate production (aerobic glycolysis), during expansion of healthy donor T cells in a perfusion-based microbioreactor^37^. Here, we perform repeated culture runs with destructive measurement of gold-standard attributes of cellular metabolism, to validate that our model is capable of estimating rates of glycolysis and OxPhos. As CAR T expansion dynamics are known to be affected by the nature of the expansion platform^38^, our tool has been designed to accommodate different reactor types and operating modes such as batch, perfusion, and fed batch cultures, and for both cases where continuous data is available through online sensors, and for cases with intermittent manual measurements. Applying our model to both healthy donor and patient-derived cells suggests the potential for using early metabolic indicators to predict final cellular attributes.

## Results

### Computational modelling allows estimation of key metabolic parameters from continuous bioreactor measurements

To estimate parameters of growth and metabolism of cells growing in a bioreactor, we developed a software tool that has at its core an Ordinary Differential Equation (ODE) model (Figure 2, See methods). Specifically, the tool performs optimization of the ODE-model to estimate rate parameters that minimize differences between model predictions and experimental measurements (online and offline). The model takes as input continuous or infrequent measures of cell density, cell count, dissolved oxygen concentrations and metabolite concentrations. The model then provides as output the required rates of growth and metabolism, such as oxygen consumption rates and lactate production rates which can be validated through experimental assays. Further details can be found in the Methods.

**Figure 2.**
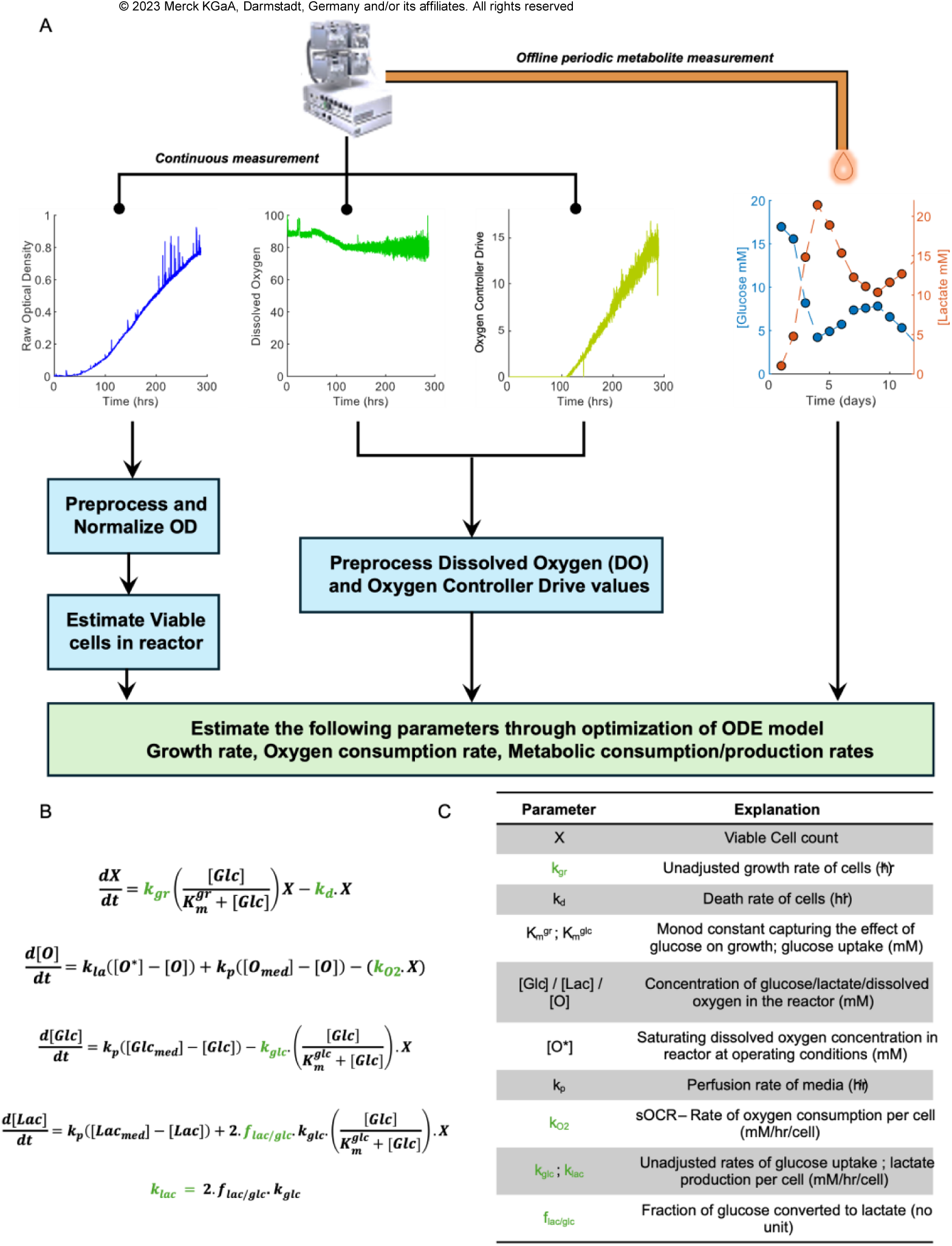
A modelling tool to estimate growth and metabolic rates of cells cultured in a microbioreactor. (A) Schematic illustrating the inputs, outputs and internal processing involved in estimating the rates of growth and metabolism for cells growing inside a microbioreactor. The bioreactor is equipped with online sensors that continuously measure optical density (OD), dissolved oxygen (DO) and relative supply fractions of air and pure oxygen. In addition, we also perform manual offline measurement of daily metabolite concentrations. These are the inputs to our tool that uses ordinary differential equations (ODEs) and optimization to estimate rates of growth, oxygen consumption and metabolic rates such as glucose consumption and lactate production per cell while accounting for other factors such as change in cell count and perfusion rates (or media exchange schedules for batch cultures). (B) The system of ODEs used by our tool to estimate the target growth and metabolic rates (indicated in green). (C) Table listing the variables and parameters in the ODE model and what they represent.

We first applied our tool to analyze CAR T cell expansion in a continuous perfusion-based microbioreactor. Details of the experimental design and setup can be found in the Methods section and in Supplementary Figure 1. The bioreactor is equipped with online sensors that provide real-time continuous measurements of optical density (OD, a measure of cell density), dissolved oxygen concentration (DO) and the proportion of pure oxygen-vs-air supplied to the reactor to maintain DO at a setpoint (Oxygen controller drive). In addition to these online measurements, we also performed daily offline measurements of glucose and lactate concentrations in the perfusate (Figure 2A). The combination of real time data from the reactor and daily offline measurements provides a rich dataset as input to the modelling. For this data-rich case, our tool uses a simplified ODE model (ignoring ODEs for glucose-dependent cell division and death), as the available measurements of real-time optical density can be used to reasonably approximate the viable cell count (VCC) inside the reactor at any point of time (Figure 2B-C and Methods).

Using the online OD and DO values and offline metabolite measurements as input, the tool estimates the rate of cellular growth, specific oxygen consumption rate (sOCR = mean OCR per cell), and rates of glucose consumption and lactate production per cell. We applied the model to three expansion experiments, with each experiment using commercially available T cells from a different healthy donor (n = 3 biological replicates with one run each). Briefly, healthy T cells were inoculated in the microbioreactor on Day 0 and activated using Dynabeads. CAR-T constructs were introduced into these cells via lentiviral transduction on Day 1, and the cells were grown in the reactor until harvested on Day 12. Figure 3A shows the continuous estimates of sOCR obtained when applying our tool to the three expansion runs. We observed an increase in modelling-estimated sOCR between Days 1-2, followed by a gradual decrease until day 6, after which it remains steady until Day 12 (day of harvest). This is in alignment with earlier studies that show an increase in OxPhos following activation of T cells^14,17–19^. Figure 3B shows the amount of lactate produced per cell for each of the three donors. Note that unlike the sOCR estimation, the estimated lactate production rates are not continuous, but an average rate over the interval between two successive metabolite measurements (in our case, daily). Similar to the sOCR profile, lactate rates show an increase until day 3 and deceleration between days 3-6, after which they remain steady until Day 12. A similar plot for glucose consumption per cell can be found in Supplementary Figure 2. The observed metabolic trends also agree with previously known T cell proliferation kinetics, where growth rate peaks immediately following activation, followed by deceleration and plateau phases. Under normoxic conditions, only a part of glucose consumed by a cell is exported as lactate into the media following glycolysis (aerobic glycolysis). The rest of the glucose intake is diverted internally to produce energy via pyruvate and TCA cycle, or towards anabolism. The modelling-derived estimates of glucose consumed per cell and lactate produced per cell were used to compute the fraction of glucose used by a cell to produce lactate (f_lac/glc_), which is a measure of aerobic glycolysis (Figure 3C). Similar to raw consumption/production rates, the levels of aerobic glycolysis increase following activation and show a steady decrease until Days 11-12.

**Figure 3.**
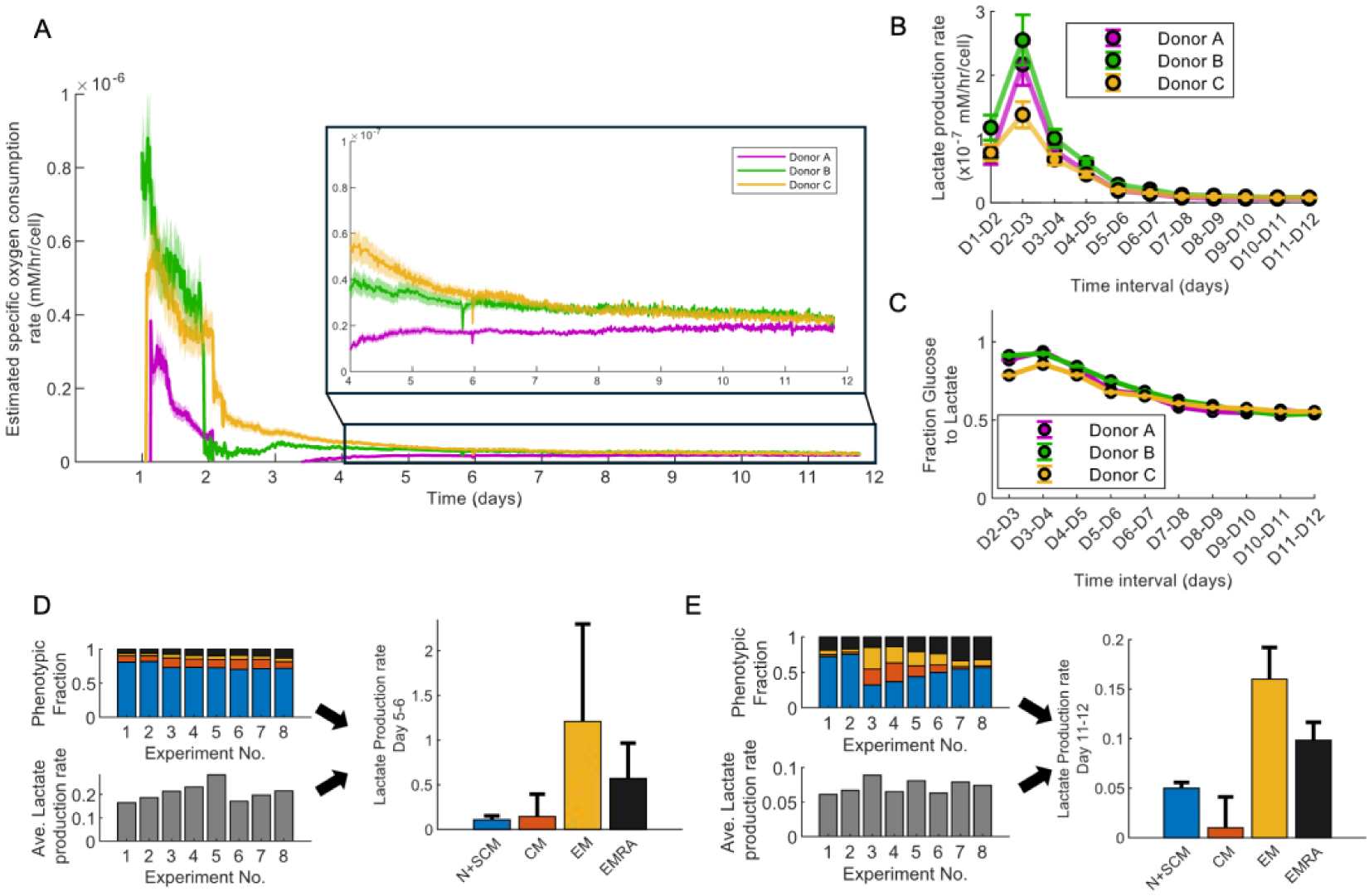
Modelling estimates rates of oxygen intake and glycolysis of healthy CAR T cells. (A) Continuous estimates of specific oxygen consumption rates (sOCR) for three commercially available healthy T cells expanded in the microbioreactor following activation and CAR transduction (n = 1 per sample). Dark lines indicate the median sOCR for each of the T cell samples and the shaded region indicates median absolute deviation, the uncertainty arising from estimating viable cell count in the reactor using OD values (See methods for more details about VCC estimation). At both early stages (Days 1-4) and late stages of the expansion process (Days 10-12), subtle differences can be observed between the estimated sOCR values for the three samples. Inset shows zoomed in plot for days 4-12. (B) Lactate production rates (adjusted) for the three healthy CAR T cell samples. Unlike sOCR estimates, the lactate production rate is an average rate across a 24-hour interval, as it is based on daily offline measurements of metabolite concentrations. Error bars indicate the uncertainty arising from VCC estimation from OD values (mean ± SD). Differences can be observed between the three samples during the early days (Days 1-4) immediately following T cell activation. (C) Fraction of glucose intake that is used for lactate production per cell, for the three healthy CAR T cell donor samples (mean ± SD). This is estimated from the rate of glucose uptake (Supplementary Figure 2) and the rate of lactate production per cell and is a measure of aerobic glycolysis performed by the cells. In all cases, the cells are close to being 100% glycolytic following activation (between days 3-4), following which there is a steady decline in the levels of aerobic glycolysis of cells in the reactor. (D) The day 6 offline flow cytometry measurement of phenotypic fractions (top left) and the estimated overall average lactate production rate (mM/hr/cell) for the interval D5-D6 (bottom left) are used to estimate phenotype-specific lactate production rates (right). The crude phenotype-specific estimates on the right show qualitative agreement with known T cell phenotype specific metabolic trends. (E) Same as (D) but obtained using day 12 offline phenotypic measurements and average lactate consumption rates for the interval D11-D12.

While the rate estimates from modelling are per cell, they represent average rates across the entire population in the reactor. They do not account for the fact that the reactor contains T cells of different differentiation phenotypes which can have different metabolic profiles. We performed flow cytometry on Days 6 and Days 12 of the expansion process to estimate the relative proportion of the four major T cell phenotypes based on their expression of the surface markers CD45RA and CCR7, separating them into CD45RA^+^CCR7^+^ Naïve + Stem Cell Memory cells; CD45RA^−^CCR7^+^ Central memory cells; CD45RA^−^CCR7^−^ Effector memory cells; and CD45RA^+^CCR7^−^ Terminally differentiated effector cells^37^. Using these relative phenotypic proportions and the average estimated lactate production rates on Days 6 and 12 as input, we used a simple linear regression model to estimate phenotype-specific lactate production rates for Day 6 (Figure 3D) and Day 12 (Figure 3E). Metabolic rates of individual phenotypes in a heterogeneous population may otherwise be impossible to ascertain through experimental measurements, as purifying subtypes could also alter their metabolism. Thus, the computational method we use might provide the only avenue to estimate phenotype-specific metabolic rates in a non-destructive, sterile manner. Figures 3D-E show relative lactate production rates by each phenotype (estimated), which is in concordance with previous studies of T cell differentiation and metabolism discussed earlier. In brief, Naïve+SCM (T_N_+T_SCM_) cells have very low overall glycolysis, Effector memory (T_EM_) cells are predominantly glycolytic in nature, and T_EMRA_ cells experience a drop in metabolism compared to T_EM_ cells. Central memory (T_CM_) cells are known to be more OxPhos dependent and thus show low to no aerobic glycolysis. The same relative patterns are recapitulated by the estimated phenotype-specific rates on both Days 6-12. Thus, modelling provides deeper insight into metabolic activity within a cell population than might be apparent from looking at changes in absolute metabolite concentrations in media. While the current analysis uses model-estimated metabolic rates and flow cytometry-measured phenotypic fractions to compute phenotype-specific rates, the analysis can also be reversed to estimate phenotypic fractions from overall metabolic rates without requiring sampling and flow cytometry measurements, as more quantitative data becomes available on phenotype-specific rates in different contexts.

### Model-derived estimates of glycolysis and oxygen consumption are in concordance with experimental measurements

To estimate the reliability of our model-estimated parameters, we performed new CAR T expansion runs comparing model-estimated values against experimentally measured values of respiration (oxygen consumption rate, OCR, and extracellular acidification rate, ECAR) and glycolysis (proton efflux rate, PER) using the Seahorse XF analyzer. For all runs, T cells were seeded into the microbioreactor on Day 0 and activated. For metabolic model validation experiments, T cell were seeded and activated, but not transduced with a CAR transgene (as per usual protocol in Supplementary Figure 1), as our goal was only to compare modelling-estimates of metabolic rates vs assay-estimates. Experiments were performed in two conditions – native, or in the presence of 2-Deoxy-D-glucose (2DG), a known inhibitor of glycolysis and overall metabolism. For each condition, we performed three expansion runs - all with the same experimental conditions and protocols, but terminated and harvested on different days (Day 2, 3 or 7) (Figure 4A). For the run terminated on day 7, an additional intermediate cell sampling step was performed on Day 4, by removing 50uL of cells from the reactor. For every run, metabolite concentration measurements were typically performed once or twice a day, except on the day of final harvest/sampling. On harvest/sampling days, we ensured two mandatory metabolite measurements were performed within 3-hrs of each other, with the second measurement being just prior to the final harvest/sampling. This allowed us to compute average lactate production rates corresponding to a 3-hr window just prior to extracting cells for metabolic estimation on the Seahorse analyzer (assay-estimate).

**Figure 4.**
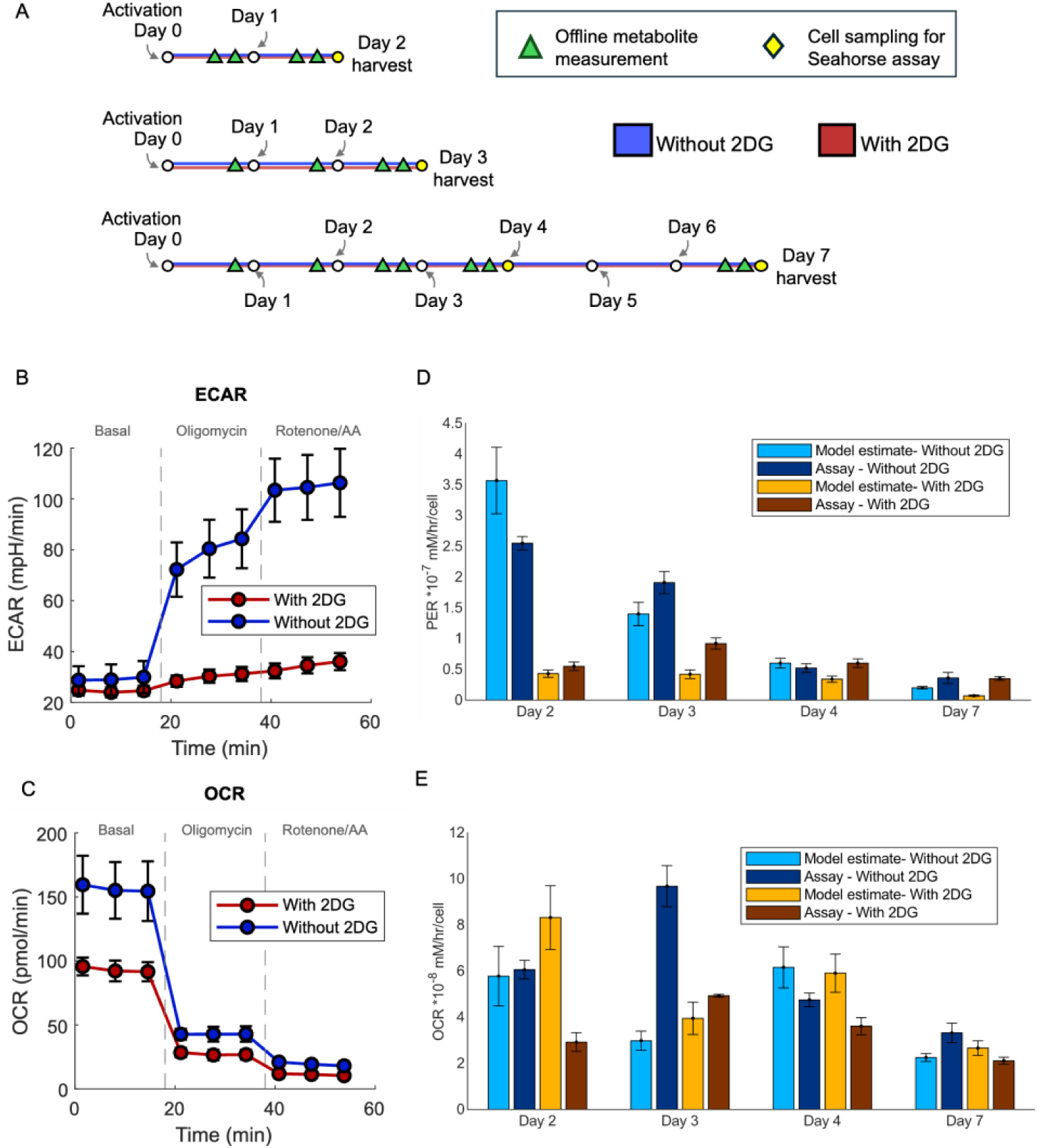
Independent measurements of metabolic rates using the Seahorse assay validate model-estimated metabolic rate parameters. (A) Schematic illustrating the design of our experiments to estimate the rates of oxygen consumption and lactate production (assay-estimate) using the Seahorse Extracellular Flux (XF) analyzer. Briefly, we performed three independent expansion runs of healthy T cells (not CAR transduced) grown in the microbioreactor, with identical conditions (inoculated on Day 0). The runs were terminated and cells harvested on different days for each run (Day 2, 3 and 7). For the 7-day run, there was an intermediate cell sample taken on day 4. For the harvested and intermediate cell samples, assay-estimates for Oxygen Consumption Rates (OCR) and Extracellular Acidification Rates (ECAR) were obtained using the Seahorse analyzer. For each run, the start of a new day is indicated by a white circle, and the number of metabolite measurements performed each day is indicated as green triangles. On the day of harvest (or intermediate sampling), two metabolite measurements were performed roughly 3 hours apart just preceding cell collection. This entire setup was repeated in a second condition where the cells were exposed to a metabolic inhibitor 2-Deoxy-D-glucose (2DG). (B-C) Example plots of OCR and ECAR obtained from the Seahorse analyzer for one cell sample (7-day run) in both native and 2-DG conditions (mean ± SD). (D) Comparison of model-estimated (pale) and assay-estimated (dark) rates of lactate production, for the native case (blue) and the 2DG case (orange) at all four points of cell collections (Days 2, 3, 4 and 7). Error bars for the modelling-estimates arise from uncertainties in VCC estimation (mean ± SD), whereas those for assay-estimates arise from technical replicates (n = 3) for the Seahorse assay for each cell sample (mean ± SD). (E) Comparison of model-estimated and assay-estimated values of OCR (mean ± SD) per cell similar to (D).

Figures 4B and 4C show sample plots for extracellular acidification rate (ECAR) and oxygen consumption rate (OCR) obtained for the Day 7 sample using the Seahorse metabolic assay. Similar plots for samples from days 2, 3, and 4 can be found in Supplementary Figure 3. ECAR values were further processed to obtain Proton efflux rates (PER) which is a measure of protons generated by cells into the media. PER was further broken down into glycolytic PER and mitochondrial PER. Glycolytic PER (GlycoPER) refers to protons released specifically from lactic acid and is thus directly comparable to lactate production rates from modelling. Figure 4D shows the modelling-estimated and Seahorse assay-estimated lactate production rates for both the native and 2DG conditions for cells sampled on days 2, 3, 4 and 7. It can be seen that there is very good agreement on both absolute and relative scales between the modelling-estimated and assay-estimated rates, with a RMSE of 0.463*10^−7^ mM/hr/cell for GlycoPER estimates and 3.23*10^−8^ mM/hr/cell for the sOCR estimates. There is an overall monotonic decrease in lactate production rates from Day 2 to Day 7, and in all cases the 2DG condition is lower or comparable to the native condition. While the estimated lactate production rates for the final harvest/cell sampling timepoint are shown in Figure 4D, intermediate lactate production rates for each run can be seen In Supplementary Figure 4. Similar to Figure 4D, Figure 4E shows final estimated values for the specific oxygen consumption rates. Except in a couple of cases in earlier data (Day 2 with 2DG, Day 3 without 2DG), the modelling estimates are consistent with assay-estimated rates. The fact that modelling predictions vary in either direction (2.85-fold overestimation in the Day 2 case and 3.25-fold underestimation in the Day 3 case) and affect both conditions (with 2DG for the Day 2 case and without 2DG for the Day 3 case) suggests that there is no systematic bias in the model estimated rates. Instead, these differences are possible consequences of random errors in cell count measurements and DO measurements in the first few days and can be eliminated with further data. Taken together, these results serve as validation for metabolic rate parameters estimated via our tool.

### The model can be successfully applied without continuous data measurements, and to non-perfusion reactor systems without online measurement capabilities

Results in the previous sections were obtained using the data-rich case of a perfusion-based microbioreactor with online sensors. To understand model performance when continuous data is not available, we synthesized pseudo-datasets by downsampling the online sensor data to one cell count “measurement” per day. As shown in Figure 5A, the raw OD values were converted to estimates of viable cell count (VCC), from which we sampled points at 24-hour intervals, to mimic the process of daily offline measurements of cell counts (similar to the metabolic measurements). We then used this as input to the model, which now also includes ODEs to model cell growth and cell death, to estimate VCC at any point of time between offline measurements (Methods). We call this the downsampled microbioreactor (MBR) dataset. Figure 5B shows the result of applying the full model to this downsampled MBR dataset for all the Seahorse validation runs. The model-estimated lactate production rates show overall reasonable concordance (RMSE = 0.532*10^−7^ mM/hr/cell), but greater individual deviations from the corresponding assay-estimated PER values, than in the case of using the full dataset. While the largest differences when using the full dataset occur during Day 2 where the absolute values are the highest, the largest differences when using the downsampled dataset occur during Day 4, when the absolute values are low and noise from cell count measurements are likely low. This implies that the quality of rate estimation from downsampled dataset is much poorer compared with the full dataset, than implied by the RMSE values. This serves to illustrate two points. First, continuous measurements of cell count can provide much better estimates of interval metabolic rates than daily cell count measurements. Second, in the absence of continuous measurements, the model is still capable of obtaining crude estimates of metabolic rates of cells within the reactor (with frequent estimates >3-fold difference from assay measurements). Figure 5C shows how each of the six underlying parameters of the full ODE model (see Methods and Figure 2) vary significantly over time to provide the best fit between the model-estimated and Seahorse measured rates for one specific run (terminated on Day 7) with and without 2DG. Similar plots for the runs terminated on days 2 and 3 can be found in Supplementary Figure 5.

**Figure 5.**
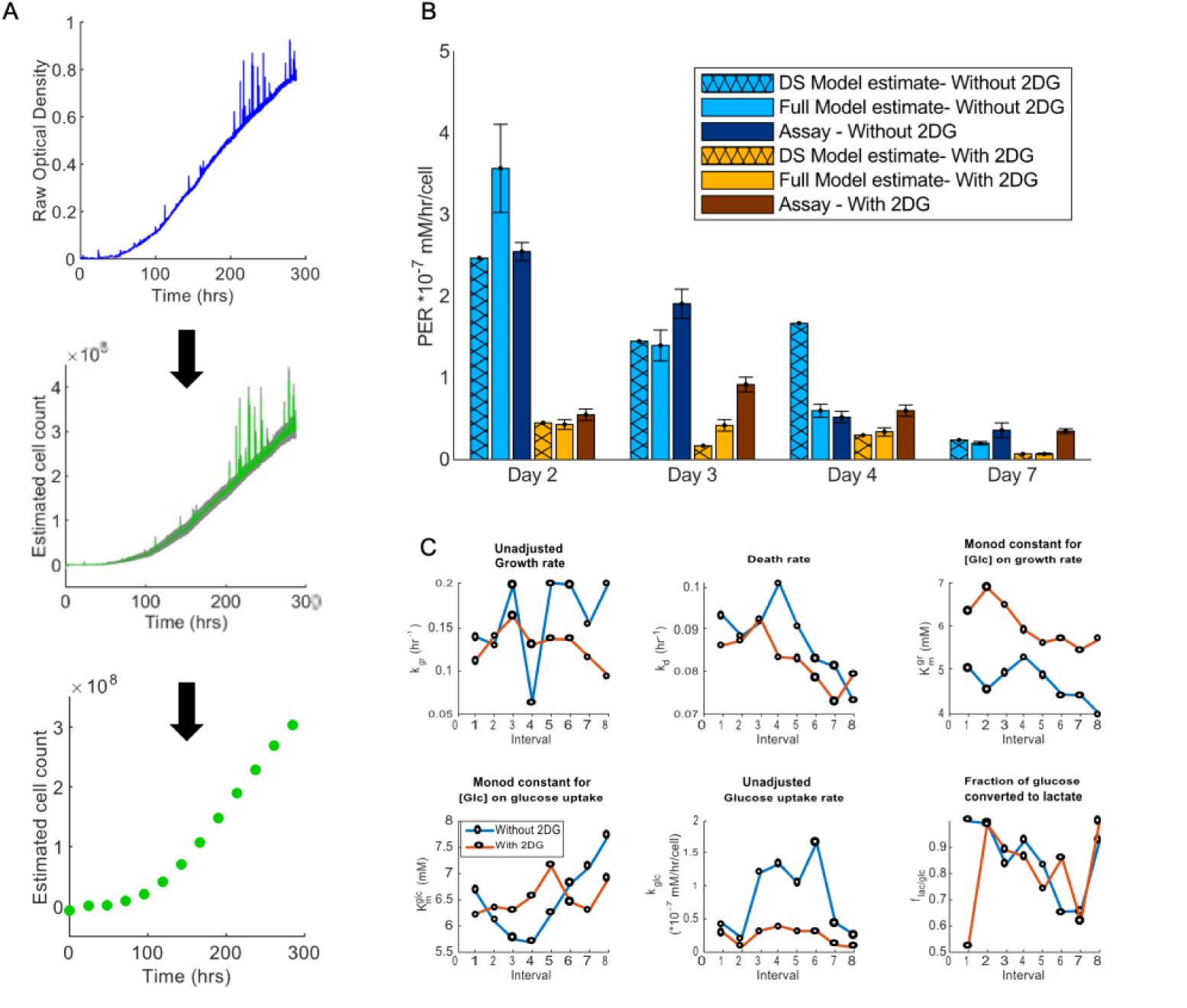
Application of modelling tool to cell cultures without continuous online measurements. (A) Workflow illustrating the process of obtaining the downsampled MBR dataset from the full set of microbioreactor data. The continuous OD data (blue) is converted to estimated viable cell counts (VCC, continuous green line) using techniques outlined in the methods and Supplement. From this we sample VCC values 24 hours apart to simulate the process of obtaining a daily offline cell count measurement. (B) Comparison of model-estimated rates for the full dataset (pale), downsampled dataset (hatched pale) and assay-estimated (dark) rates of lactate production, for the native case (blue) and the 2DG case (orange) at all four points of cell collections (Days 2, 3, 4 and 7). Error bars for the modelling-estimates arise from uncertainties in VCC estimation (mean ± SD), whereas those for assay-estimates arise from technical replicates (n = 3) for the Seahorse assay for each cell sample (mean ± SD). (C) Best-fit estimates for the daily average of six rate parameters/variables from the full ODE model (refer to Figure 2C), for both the native (blue) and 2DG (red) conditions. For illustration, we only display the estimates for the 7-day run, showing significant variability between estimates for successive days.

To demonstrate generality of our approach, we also applied our tool to a non-perfusion bioreactor system commonly used for clinical CAR-T cell manufacturing – a gas permeable G-Rex 24 well plate (8ml volume). Supplementary Figure 6 shows the experimental design, media exchange schedule and measurement frequencies for the G-Rex cell culture experiments. For these experiments, offline cell count measurements were performed only on days 0, 1, 6 and 12. As with the downsampled MBR dataset, we applied our tool to the G-Rex dataset to estimate metabolic and growth rates in the absence of continuous cell count measurements. Supplementary Figures 7-8 shows the rate estimates obtained when using the offline cell count measurements as is, or when interpolating between actual measurements to mimic daily cell count measurement respectively. In both cases, the model provides good fit between the measurements and predictions, but the underlying rate parameters show greater variability than obtained using continuous measurements, similar to the case of the downsampled MBR dataset. For comparison, we re-analyzed the downsampled MBR dataset, this time interpolating with increasing frequency to mimic offline VCC and metabolite measurements *n* times a day (*n* ∈ {1, 2, 3, 4, 8}). For the single run tested, we observed that obtaining 2 or more measurements a day yielded rate parameters very similar to those obtained using continuous OD data from the full MBR dataset (Supplementary figure 9), even though the downsampled data analysis includes more ODE equations for growth kinetics.

**Figure 6.**
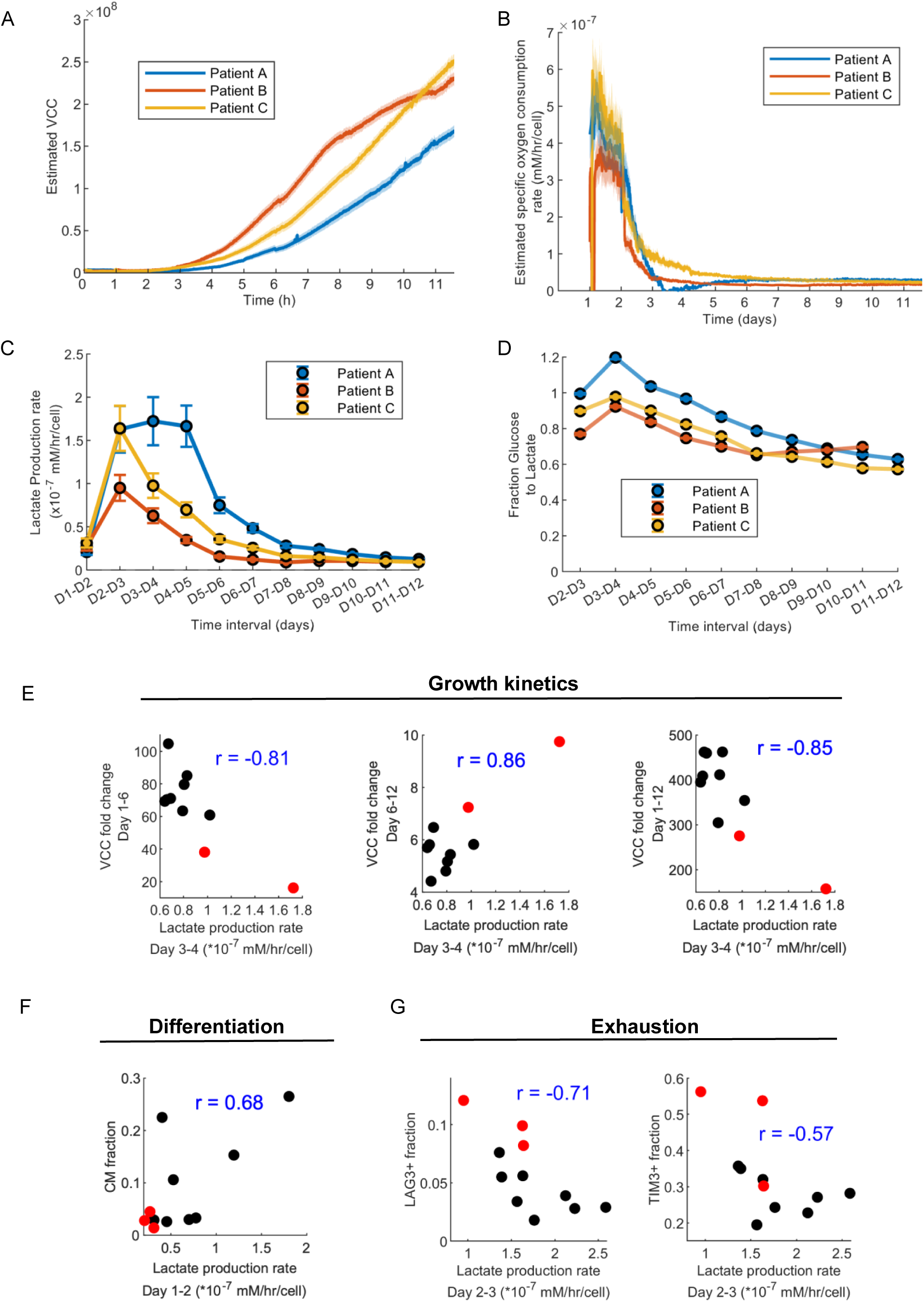
Application of modelling tool to cultures of patient-derived T cells shows early metabolic indicators correlate with final product attributes. (A) Estimated VCCs during the expansion of three different patient-derived T cells in the microbioreactor (n = 1), following activation and CAR transduction. VCCs were estimated from continuous OD values as outlined in Supplement. Dark lines represent the mean values and shaded regions represent the 10^th^ and 90^th^ percentile values of VCCs estimated from the OD. (B) Continuous estimates of specific oxygen consumption rates (sOCR) for the three patient cell expansions. Dark lines indicate the median sOCR for each of the T cell samples and the shaded region indicates median absolute deviation, the uncertainty arising from estimating viable cell count in the reactor from the recorded OD values. (C) Lactate production rates for the three patient samples. Unlike sOCR estimates, the lactate production rate is an average rate across a 24-hour interval, as it is based on daily offline measurements of metabolite concentrations. Error bars indicate the uncertainty arising from VCC estimation from OD values (mean ± SD). (D) Fraction of glucose intake that is used for lactate production per cell for the three patient samples (mean ± SD). This is estimated from the rate of glucose uptake and the rate of lactate production per cell and is a measure of aerobic glycolysis performed by the cells. (E) Scatter plot of lactate production between days 3-4 and growth parameters – fold change in estimated VCC between days 1-6, between days 6-12 and between days 1-12. Black circles indicate expansion runs with healthy donor cells and red circles indicate expansion runs with patient cells (n = 2 as a technical issue prevented offline cell count measurement for patient B on Day 12). Text in panel shows the value of Pearson correlation coefficient (r). Interestingly, Lactate production rates displays opposing relationships with growth rates during early (days 1-6) and late (days 6-12) phases of expansion. (F) Scatter plot showing rate of lactate production between days 1-2 and the fraction of central memory (T_CM_) type T cells on day 12. (G) Scatter plot of lactate production rate between days 2-3 and exhaustion markers – fraction of LAG3+ cells and fraction of TIM3+ cells.

### Application of model to expansion of patient-derived cells shows potential relationships between early metabolism and final cellular outcomes

Having demonstrated the successful application of our tool to the expansion of CAR-T cells from healthy donors, we proceeded to apply it to CAR-T cells manufactured from material obtained from cancer patients to better simulate clinical CAR-T production. Briefly, peripheral blood mononuclear cells (PBMCs) were extracted from discarded leukapheresis tube sets for three adult lymphoma patients (anonymized). T cells were isolated from the PBMCs and 2 million T cells from each patient were inoculated in the microbioreactor, following the same 12-day activation, transduction and expansion protocol as with the healthy donor cells (See methods). Figure 6A shows the estimated viable cell count from the online OD measurements for the three patients. The patient cell expansion process resulted in Day 12 viable cell counts that were 30-50% lower than the healthy donor cells, although inoculated with the same 2 million cells on Day 0. In addition, patient cells showed significant differences between each other in growth kinetics (Figure 6A) and final cellular count on harvest. Figure 6B-D show estimated rates of oxygen consumption, lactate production and glucose-to-lactate fractions for the three patients. The estimated rates from each patient show clear differences in the early few days. This demonstrates inter-patient variability in T cell phenotypes and growth/differentiation kinetics that confound standardization of the CAR T cell manufacturing process. Understanding how these differences in metabolism relate with CQAs of final product can be instructive for process optimization and standardization of CAR T manufacture.

### Early metabolic parameters correlate with CQAs of harvested cellular product

Prior studies have suggested that early metabolism and growth kinetics following activation can affect final cell viability, proliferation capacity, phenotypes and exhaustion. We hypothesized that early rate estimates obtained using modelling (specifically lactate production rates) could provide indications toward final cellular output. To explore this possibility, we collated data on growth kinetics, differentiation status and exhaustion metrics for eight healthy-donor runs and the three patient runs using the perfusion microbioreactor. For growth kinetics, we used pairwise fold changes between offline cell count measurements on days 1, 6 and 12. Differentiation and exhaustion were estimated using flow cytometry performed on Days 6 and 12. For differentiation, the relative expressions of surface markers CD45RA and CCR7 were used to classify cells as described earlier. For exhaustion, we measured the relative expression of known exhaustion markers LAG3, TIM-3 and PD-1. We then studied the correlations between early lactate production estimates (Day 1-2, Day 2-3, Day 3-4 and average of Day 1-4) against all measured growth, differentiation and exhaustion metrics. Figure 6E-G shows only those relationships between early lactate production and final cellular metrics, that both showed high Pearson correlation coefficient (|r| > 0.5), and passed our statistical significance threshold of p<0.05 for non-zero r (corrected for multiple hypothesis testing). In each panel, black circles represent the healthy donors, and red circles represent patient cells. Interestingly, Figure 6E shows that the estimated lactate production rate between days 3-4 (L_3-4_) appears to have contrasting relationships with cell growth during the first half and the second half of the expansion process. L_3-4_ appears negatively correlated with fold change of VCC between days 1-6 (FC_1-6_) but is positively correlated with the fold change of VCC between days 6 and 12 (FC_6-12_). This raises questions about whether glycolysis may have different effects on the cellular expansion process immediately following activation and later once differentiation and exhaustion patterns are more firmly established. Overall, higher L_3-4_ appears to suggest lower FC_1-12_ i.e., lower overall proliferation during the entire 12-day expansion period.

Figure 6F shows that very early lactate production rate between Days 1 and 2 (L_1-2_) appears to be correlated positively with the proportion of T_CM_ cells on Day 12. Figure 6G shows that Lactate production rates between days 2 and 3 (L_2-3_) are negatively correlated with expression of exhaustion markers LAG3 and TIM-3. This is along the lines of reports from other disease contexts such as Type 1 Diabetes, where suppressing glycolysis was found to induce exhaustion in T cells^39^. Correlations uncovered using such techniques are not necessarily an indication of physiological cause-and-effect, but rather provide relationships that can be studied in greater detail using targeted experimental design. Their ability to predict final CQAs using a simple regression model is shown in Supplementary Figure 10. Overall, our work aims to illustrate the value in studying early metabolism of CAR T cells in order to predict critical quality metrics of the final cell therapy output.

## Discussion

Patient-specific treatments such as autologous cell therapy are increasingly preferred in many refractory cases of cancer where generic treatments might fail for varied reasons. However for autologous therapies, the source material (patient T cells) can be very variable for each patient due to both intrinsic heterogeneity and prior treatment history. For such variable cases, a targeted and adaptive manufacturing process is desirable for the following reasons. First, they can decrease production times which is preferred, as shortening vein-to-vein time for potent formulations such as CAR-T therapies by just days can result in increased life expectancy (in years) and quality of life^40,41^, translates to lower manufacturing costs, and frees up critical manufacturing infrastructure for higher throughput. This has also been borne out by an increasing focus on accelerating CAR T manufacturing from the typical weeks^11,42–44^ to a few days^44–50^. Second, an adaptive process may decrease the probability of CAR T manufacturing failure (which can reach up to 13%^51^) by identifying early irregularities and their root causes, and rescuing the process from a complete failure to grow cells, or out-of-specification products. Third, adaptive processes can help improve therapeutic efficiency and potency by improving non-mandated key cellular attributes such as phenotypes, exhaustion and proliferation capacity. We posit that cellular metabolism provides a good handle for developing such adaptive processes and that knowledge of metabolic states of cells in the bioreactor may allow for early identification of key cellular attributes through the relative activity of OxPhos and aerobic glycolysis. In this work, we have developed a tool for such metabolic state estimation using computational modelling.

For T cell therapies, computational modelling has typically focused on protein design for CAR sequences ^52,53^, modelling the dynamics of infused CAR T cells^54^, studying patient responses^55^, CAR T – tumour interactions and dosing^56^, CAR T pharmacodynamics^57^, or to minimize the cost of CAR T manufacturing by optimizing media supply to a bioreactor^58^. Our work is one of the first attempts to apply modelling for state estimation in cell therapy manufacturing. We first present measurements and integrated modelling that allow for continuous estimation of metabolic state for the cells in culture. We then provide model validation in the form of experimentally measured metabolic rates from a metabolic analyzer (Agilent Seahorse) assay, and show the potential for using metabolic rates to estimate real time phenotypic fractions in culture. We also demonstrate applicability of our tool to T cells obtained from patients, characterizing biological variation inherent in patient-derived cells. Most important of all, using both healthy donor and patient-derived T cells, we provide the first proof-of-principle to show that early metabolic indicators such as lactate production rates are correlated (|r| > 0.5) with key cellular attributes of the final output such as proliferation capacity, differentiation (T_CM_ fraction) and exhaustion (fraction of LAG3+ and TIM3+ cells), an idea that has not been previously reported in literature. A notable feature of our work is the ability to estimate metabolic states in a sterile manner which is an important requirement of clinical GMP-certified manufacturing processes. While most results in this work have been obtained using in-house CAR T cell expansion data from a perfusion based microfluidic bioreactor with 2mL working volume^37^, our model can also be applied to different types of reactor systems (batch/fed-batch/perfusion) with or without continuous online sensor data.

While this study demonstrates our model’s capabilities to recapitulate experimental measurements, estimated metabolic parameters can also be sensitive to the accuracy of cell count inference or metabolite measurements (and their derivatives). Additionally, the correlation results we present here are based on a small number of expansion runs performed using healthy donor T cells and patient-derived cells, and primarily show proof-of-principle without implying cause-and-effect relationships. Further data to establish causal relationships between early indicators and final CQAs can be generated using a more detailed and systematic experimental design developed using the Design-of-Experiments or systems modelling approach coupled with more frequent offline measurements of additional metabolites such as glutamate/glutamine, pyruvate, serine etc.

Frequent measurements of metabolites would also improve the accuracy of modelling in estimating metabolic rates. As we demonstrate for cases without online sensors for continuous measurements (e.g., G-Rex systems), the high variability we observed between successive estimates of some estimated rate constants (e.g., growth rate, death rate and fraction of glucose to lactate) go against conventional physiological wisdom. This illustrates a caveat of the model when there is limited measured data – that mathematical optimal fit may not always conform to physiological behaviour. In the absence of more frequent or continuous measurements, the model becomes an underdetermined system that can be satisfied by multiple parameter combinations to similar degrees, providing good match between modelling and experimental measurements. With more constraints (in the form of frequent/continuous measurements of cell counts or metabolites), uncertainty in the parameter space can be reduced to provide more physiologically meaningful estimates of metabolic rate constants. While not very common, continuous measurements of metabolite concentrations and cell densities have been demonstrated in other studies, using Process Analytical Technologies (PATs) such as Raman spectroscopy^59^. Integrating such technologies for real time monitoring would help to refine such models and also generate more data, allowing for the use of more powerful models such as neural networks for state estimation. Thus, real-time data collection, regardless of bioreactor type, could be an excellent goal for next generation reactor and process design.

The topic of adaptive processes for phenotype enrichment also requires further study. Maximizing the yield of certain phenotypes (e.g., T_CM_) may not necessarily imply maintaining an enriched/purified phenotype population for the entirety of the culture, as phenotype-phenotype communication shapes metabolism and growth dynamics of individual subtypes. Bioreactors with perfusion and rapid mixing deprive cells of paracrine and transient juxtacrine (contact-based) signals that likely affect cell fate.. While estimating phenotypic fractions from metabolic rates is extremely useful from an adaptive process point of view, there is yet little knowledge of how phenotype-specific metabolic rates might vary under different conditions or between patients. Further studies on phenotype-specific metabolism can augment our understanding of such processes. In addition, as more clinical release standards (beyond just purity and sterility^60,61^) are defined in the future for attributes such as relative fractions of CM and naïve/SCM populations, CD4^+^/CD8^+^ ratios, cytokine profiles or potency requirements, metabolic changes associated with such attributes can be incorporated in a model predictive controller framework for adaptive process control. Overall, our tool provides one of the earliest steps towards this goal of targeted cell therapy manufacturing with shorter turnaround times and better therapeutic efficacies. With better release metrics and standards, and mapping of such standards to measurable metabolic fingerprints, such modelling tools can serve as the building blocks for adaptive process control and optimization for improving critical quality attributes (CQAs) of cell therapy products.

## Methods

### Lentivirus preparation

Lentiviral vectors encoding anti-CD19 CAR were produced, concentrated, and titrated according to the protocol described in our earlier study^37^. Briefly, packaging and transfer plasmids were transfected into HEK293T cells, lentiviral vector supernatant harvested after 48-96 h, concentrated by ultracentrifugation, and titrated using Jurkat cells.

### Preparation and operation of microbioreactor

Setup of the perfusion microbioreactor and initial priming/calibration of the microfluidic cassettes were performed according to the procedure described in our earlier study^37^. A 2 mL cell–bead mixture was injected into the growth chamber by vacuum inoculation. Transduction was performed by removing 1 ml of cell-free medium through the perfusion output, and injecting 1 ml of fresh medium containing diluted LVV into the growth chamber by vacuum inoculation. The microbioreactor cultures were controlled at the following set points: temperature of 37 °C, minimum CO_2_ levels of 5%, minimum dissolved O_2_ levels of 80% air saturation and a pH of 7.40 ± 0.05. Cell samples were taken by sampling 50 µl through the sampling port into a 1.5 ml tube. Cell-free medium samples were taken by removal of ∼200 µl perfusate (accumulated over ∼2.4 h for 1 v.v.d., 1.2 h for 2 v.v.d., 48 min for 3 v.v.d. and 36 min for 4 v.v.d.) from the perfusion output using a syringe.

### Human T cell isolation, activation, transduction and expansion from healthy PBMCs

T cells were isolated from human PBMCs from healthy donors (STEMCELL Technologies) using EasySep Human T Cell Isolation Kit (STEMCELL Technologies) and resuspended in AIM V Medium (Thermo Fisher Scientific) supplemented with 2% human male AB serum (Merck Sigma-Aldrich) and 100 IU ml^−1^ recombinant human IL-2 (Miltenyi Biotec). For activation, purified T cells at 1 million cells per ml were mixed with Dynabeads Human T-Expander CD3/CD28 (Thermo Fisher Scientific) at a 1:1 cell to bead ratio, and 2 ml of cell– bead mixture (containing 2 million cells and 2 million Dynabeads) was seeded into each gas-permeable well of a G-Rex 24-well plate (Wilson Wolf) or each microbioreactor cassette (Erbi Biosystems, MilliporeSigma). One day after activation, 1 ml of cell-free medium was removed, and then 1 ml of fresh medium containing diluted LVV was added to the wells or cassettes at a MOI of 5. One day after transduction, 6 ml of fresh medium was added to achieve the final culture volume of 8 ml for the gas-permeable wells, and perfusion was started at 1 v.v.d. for the microbioreactor. For the gas-permeable wells, 6 ml medium was exchanged every other day, with a constant culture volume of 8 ml per well. For the microbioreactor, the perfusion flow rate was increased by 1 v.v.d. every other day until a maximum of 4 v.v.d., with a constant culture volume of 2 ml. T cells were expanded over 10 days, for a total process duration of 12 days.

### Human T cell isolation and CAR T cell production from patient PBMCs

PBMCs were collected from discarded leukapheresis tubing sets (Spectra Optia Apheresis System, Terumo BCT) of three anonymized adult patients with lymphoma by Ficoll–Paque density gradient centrifugation (Cytiva) for 30 min at 400 *g*. PBMCs were cryopreserved in CryoStor CS10 (STEMCELL Technologies) and then thawed for CAR T cell production following our 12 day protocol, starting with 2 million purified T cells in the microbioreactor or gas-permeable well, as described above.

### Cell count and viability

Cell counts and viabilities were measured using acridine orange (AO) and propidium iodide (PI) and/or trypan blue (TB) staining on a CellDrop Automated Cell Counter (DeNovix).

### Cell-free medium metabolite analysis

Metabolites in cell-free medium, including glucose, lactate, glutamine, glutamate and ammonia, were measured on a Cedex Bio Analyzer (Roche CustomBiotech).

### Flow cytometry

For T cell surface phenotyping, the panel used include: LIVE/DEAD Fixable Violet (Thermo Fisher Scientific), BV510 anti-CD4 (clone OKT4), BV605 anti-CD45RA (clone HI100), BV650 anti-CD8a (clone RPA-T8), PerCP/Cy5.5 anti-LAG-3 (clone 11C3C65), PE anti-CCR7 (clone G043H7), AF700 anti-PD-1 (clone EH12.2H7) and APC/Fire 750 anti-TIM-3 (clone F38-2E2) (Biolegend). Cells were washed with PBS (Thermo Fisher Scientific) then stained by incubating with LIVE/DEAD Fixable Violet stain for 20 min at room temperature. Cells were subsequently washed with FACS buffer (PBS supplemented with 2% fetal bovine serum (Thermo Fisher Scientific) and 0.1% sodium azide (Merck Sigma-Aldrich)) then stained by incubating with antibodies for 20 min at room temperature. Cells were subsequently washed with FACS buffer again before acquisition on a CytoFLEX S flow cytometer (Beckman Coulter).

### Metabolic analyzer Experiments

The reported OCR and ECAR values were first normalized to per cell values using the offline cell count obtained for each sample prior to loading onto the metabolic analyzer (Agilent Seahorse). From the normalized ECAR value, we further computed the Glycolytic proton efflux rate (glycoPER) as outlined in the Agilent documentation.

Human T cells were isolated from PBMCs (Lonza) using EasySep Human T Cell Isolation Kit (STEMCELL Technologies), and resuspended in AIM V Medium (Thermo Fisher Scientific) supplemented with 2% human male AB serum (Merck Sigma-Aldrich) and 100 IU/mL recombinant human IL-2 (Miltenyi Biotec). Purified T cells were activated with Dynabeads Human T-Expander CD3/CD28 (Thermo Fisher Scientific) at a 1:1 cell:bead ratio, and 2 mL of cell-bead mixture (containing either 2 million cells and 2 million Dynabeads for Days 3, 4, and 7 analyses, or 5 million cells and 5 million Dynabeads for Days 1 and 2 analyses) was seeded into each microbioreactor cassette (Erbi Biosystems, MilliporeSigma) in the presence or absence of 2.5 mM 2-DG (Sigma D8375). A cell sample was taken for cell count shortly after cell inoculation.

For Days 1 and 2 Seahorse analyses, media perfusion was started at 1 vvd on Day 1 post-inoculation. On each day, two perfusate samples were taken for subsequent Cedex Bio Analyzer (Roche CustomBiotech) analyses, before cell harvest. The collected cells were counted, resuspended in Seahorse XF RPMI assay medium, and then 150,000 cells were seeded in each well of a XFp PDL Miniplate for Seahorse XFp Real-Time ATP Rate Assay on a Seahorse XF HS Mino Analyzer (Agilent) according to manufacturer instructions.

For Days 3, 4 and 7 Seahorse analyses, a 1 mL media exchange to mimic transduction was performed on Day 1 post-inoculation, and then media perfusion was started at 1 vvd on Day 2 post-inoculation, 2 vvd on Day 4 post-inoculation, and 3 vvd on Day 6 post-inoculation. On each day, two perfusate samples were taken for subsequent Cedex Bio Analyzer (Roche CustomBiotech) analyses, before cell harvest. The collected cells were counted, resuspended in Seahorse XF RPMI assay medium, and then 150,000 cells were seeded in each well of a XFp PDL Miniplate for Seahorse XFp Real-Time ATP Rate Assay on a Seahorse XF HS Mino Analyzer (Agilent) according to manufacturer instructions.

### Metabolic modelling: OD preprocessing

The optical density (OD) values recorded by the microbioreactor underwent routine preprocessing before being used as input to the computational model. The necessities and methods driving individual steps of the preprocessing are described below and in Supplementary Figure 11.

Correction for reference OD: OD values are computed as the logarithm of transmission, which is the ratio of the intensities of transmitted light (I) and incident light (I_o_).

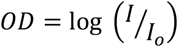

At each time step (1-2 mins), the microbioreactor measures both I and I_o_ and uses them to compute instantaneous OD values. The I_o_ is computed from a cell free section of the cassette, and is called the reference OD (OD_ref_). While the OD_ref_ values are expected to be constant through the course of the run, we noticed that in each instance, this value fluctuated (possibly due to interference from cell debris) and was generally highest at the beginning of the run. Since this created inconsistent irregularities in OD values computed at different points of time, we recomputed OD values using the measured instantaneous intensities (I) and the maximum OD_ref_ value recorded over the entire run.

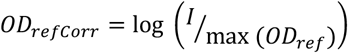

Correction for negative values: These corrected OD values were then smoothed with a moving average computed from a window of size 21 (10 minutes on either side). We also noticed that the computed ODs during the first few days of expansion had many negative OD values at early time points. This may be due to bubbles or lamination effects in the OD path during initial cassette calibration. Hence, we performed an OD correction by adding an offset as follows. For every donor/replicate, the offset value was chosen to be the smallest value (≥0) whose addition to the OD would ensure that 1% or less of corrected OD values between days 0 and 4 were negative. We call this value the *OD*_*negCorr*_.

Correction for standardization across runs: For multiple test runs of the microbioreactor, we first computed the *OD*_*negCorr*_. For each test run, we also collected the offline cell count measurements in the first 1-2 days. At this very early stage, the computed OD values are low in magnitude (typically ∼0.01), implying that in this regime, there exists a linear relationship between OD and viable cell count (VCC), as the quadratic term in the OD-VCC relationship becomes negligible. For each paired OD value and offline measurement in this regime, we computed the OD value that would correspond to 1 million cells. Comparing across all test runs, a median OD value of 0.013 was obtained for 1 million cells. Thus for each future run, a new offset was computed using the first offline cell count measurement that would result in a fixed OD value of 0.013 corresponding to 1 million cells. This offset was added to the *OD*_*negCorr*_ to give the final OD value *OD*_*final*_ that was then used for all subsequent modelling and computation.

### Metabolic modelling: estimating cell numbers from OD

Given the uncertainty in OD measurements and we sought to develop a method to estimate viable cell counts (VCCs) from OD values with a measure of uncertainty. For 38 test runs of the microbioreactor including both healthy donor and patient donor runs in this study, we collected continuous OD data and performed infrequent manual sampling of cells and measurement of offline VCCs. Pairing the measured VCCs and the *OD*_*final*_ values corresponding to the time of cell sampling, we first obtained a general quadratic fit equation using the MATLAB *fit* function. The equation was constrained to have an intercept value of 0 (reflecting an OD value of 0 for 0 cells) and positive quadratic and linear coefficients. Confidence intervals for both fitted coefficients were obtained using the *confint* command. A second series of fittings was then performed by sweeping incrementally across the confidence interval range of the quadratic coefficient, by fixing it at each value and using the *fit* and *confint* functions to obtain new confidence intervals for the linear coefficient. Repeating this process provided a *feasible area* in the linear-quadratic coefficient space whose boundaries enclosed all points that when used in a quadratic fit equation can provide a good estimate of VCC from *OD*_*final*_ values. Uniform sampling of this feasible space was performed to obtain 1000 individual points (pairs of linear and quadratic coefficients) that would serve as the final family-of-fit equations to estimate VCCS from *OD*_*final*_ with a measure of uncertainty (Supplementary Figure 12).

### Metabolic modelling: computing instantaneous specific oxygen consumption rates

First, we processed the OD data for use in inferring cell numbers. We took the microbioreactor OD data (measured roughly once per minute) and smoothed it by averaging over a sliding window of size 21 min (10 on each side of an OD value). Because the sampling periods are not exactly 1 min in each pod, we obtained synchronized readouts of all pods and all runs by interpolating the smoothed averages to obtain OD values at intervals of exactly 0.1 h (6 min), starting at 24.00 h. Each set of OD values was converted to a family of VCC curves, using the method above. Specific oxygen consumption rates (sOCR) were computed using an analytical solution to the following ordinary differential equation (ODE).

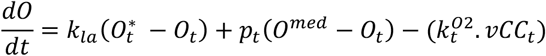

Where 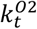 is the sOCR at time *t*; *k*_*la*_ is the volumetric mass transfer coefficient for oxygen for the microbioreactor cassette (obtained during cassette calibration); *O*_*t*_ is the dissolved oxygen concentration at time *t*; *p*_*t*_ is the perfusion rate at time *t*; *O*^*med*^ is the concentration of dissolved oxygen in fresh media, and *vCC*_*t*_ is the estimated VCC at time *t*. 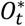is the saturating oxygen concentration at time *t* and is computed using the partial pressure of oxygen in the headspace of the reactor (based on relative fractions of pure O_2_ vs air supply in the interval *dt*). When no pure O_2_ has been supplied in the time *dt* preceding *t*, 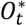 has a value of 0.203 mM corresponding to the maximum saturating oxygen concentration at a temperature of 38°C in the media. We repeated the sOCR computation with each of the 1000 fit equations (from the family-of-fits) for each microbioreactor run. This workflow yields a set of 1000 sOCR estimates for each timepoint for each run.

### Metabolic modelling: computing growth and other metabolic rates

To compute rates of growth and glucose consumption/lactate production we used the following system of ODEs

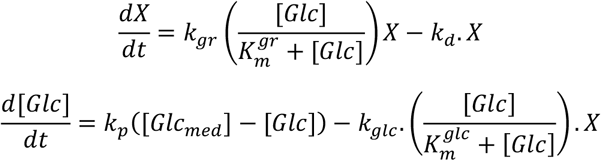

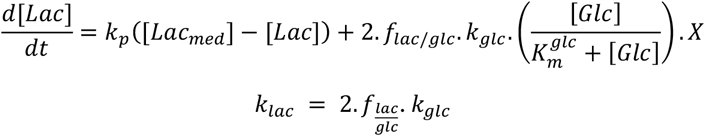

Where *X* is the VCC, *k*_*gr*_ and *k*_*d*_ are the instantaneous growth rate and death rate of cells; [*Glc*], [*Lac*] and [*Glc*_*med*_], [*Lac*_*med*_] are the concentrations of glucose, lactate in the reactor and fresh media respectively; 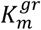 is the Monod constant reflecting the effect of glucose concentration on growth rate, *i*.*e*., it represents the concentration of glucose at which growth rate is half of the maximal growth rate; *k*_*p*_is the media perfusion rate; *k*_*glc*_ is the rate of glucose consumption per cell; 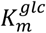 is the Monod constant reflecting the effect of glucose concentration on glucose uptake rate; *f*_*lac*/*glc*_ is the fraction of the consumed glucose that is converted to lactate. In addition, we define modified versions of glucose and lactate rates below -

Adjusted glucose consumption rate:

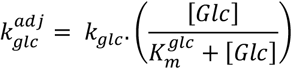

Adjusted lactate consumption rate:

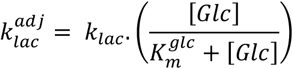

These adjusted rate parameters include the effect of instantaneous glucose concentration on the respective rates, thus reflecting net rates of glucose intake or lactate export as measured by experimental assays. Unless specifically defined as unadjusted, most results in the manuscript reflect these adjusted rate parameters.

For the case of microbioreactor where continuous measures of VCC are available from the OD, we ignore equation 1. For estimation of the unknown rates, we performed optimization to minimize the errors between measured and estimated cell counts and metabolite concentration. The optimization method was simulated annealing using the MATLAB command *simulannealbnd*, and the ODE solver used was MATLAB *ode15s*. Note that optimization steps were repeated for 100 randomly chosen fit equations (from the family of fits) for each bioreactor run, lending a measure of uncertainty to the estimated rates. To avoid local minima, we also repeated the entire optimization process five times for each fit equation, each starting with a different random seed of parameter values during initiation. This workflow yielded optimized value of metabolite rates for each time interval between successive metabolite measurements. Since in our protocol metabolite concentrations were measured once every day, the estimated rates are the average daily metabolic rates for each microbioreactor run.

### Correlation analysis

To estimate the correlation between early timepoint metabolism and final cellular attributes, we used a total of 8 healthy donor runs and 3 patient cell runs. The 8 healthy donor cells were chosen from a larger set of 11 runs obtained from pooling technical replicates from three individual donors (n = 3, 4, 4 for donors A, B and C). Prior to correlation analysis, 3 of the runs (1 from Donor A and 2 from Donor C) were discarded in an unbiased fair manner during the very first preprocessing step, as the VCCs estimated from ODs for these three runs showed significant deviation from offline cell count measurement on Day 6, implying a possible discrepancy with either offline measurement or OD-VCC fit. Correlations and associated p values were computed using MATLAB and multiple hypothesis correction was performed using the Benjamini-Hochberg method for false discovery rate.

## Contributions

N.S.J, W.-X.S., M.E.B., and R.J.R. conceived the project. W.-X.S., D.B.L.T., F.K., and Y.H. Luah conducted experiments and collected data. N.S.J., W.-X.S., and D.B.L.T., analysed data. F.L.W.I.L., M.S.-F.S. and S.Y.S. provided patient samples. Y.H. Lee, L.T.-K., M.E.B., and R.J.R. supervised the work. N.S.J., W.-X.S., M.E.B., and R.J.R. wrote the paper. All authors edited and approved the paper.

## Acknowledgements

This research is supported by Agency for Science, Technology And Research (A*STAR) under the Industry Alignment Fund - Pre-positioning Programme (IAF-PP) grant: “Assembling Screening, Productions, High-throughput Analytics, and Lentiviral Targeting for T cells (ASPHALT) (H24J4a0031)” and National Research Foundation, Prime Minister’s Office, Singapore under its Campus for Research Excellence and Technological Enterprise (CREATE) programme, through Singapore MIT Alliance for Research and Technology (SMART): Critical Analytics for Manufacturing Personalised-Medicine (CAMP) Inter-Disciplinary Research Group. The patient runs are supported by Goh Foundation Limited, Singapore to M.S.-F.S. and S.Y.S. The authors thank N. Tan at SMART CAMP, Singapore, for the coordination of patient sample collections; and K. S. Lee of Erbi Biosystems, part of MilliporeSigma, for technical assistance with the microbioreactor.

