## Supplementary figures for "Dynamic estimation of metabolic state during CAR T cell production and the relationship of early metabolism to final therapeutic product"

### Supplementary Figure 1

© 2023 Merck KGaA, Darmstadt, Germany
and/or its affiliates. All rights reserved

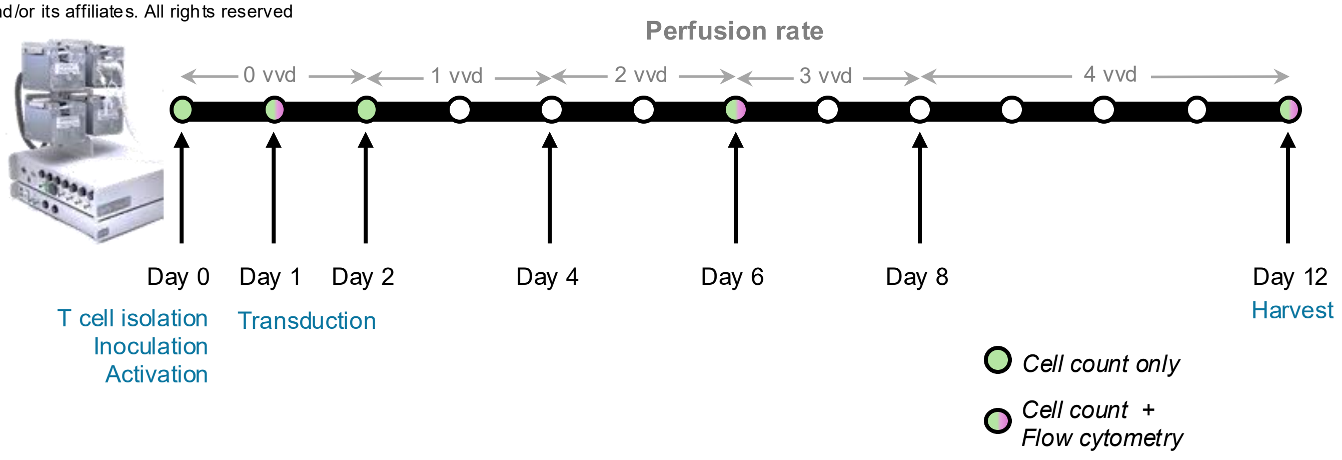


**Supplementary figure 1. Experimental design for microbioreactor manufacturing of CAR T cells from healthy donors.** PBMCs extracted from healthy donors are processed to obtain 2 million CD3+ T cells, that are inoculated into the microbioreactor on Day 0 and activated using Dynabeads. On Day 1, lentiviral transduction of CD19 CAR constructs is performed. The cells are then expanded in the microbioreactor until Day 12 when they are harvested. Perfusion rates for the reactor vary as indicated – no perfusion between days 0-2, 1 vvd (vessel volume per day) on days 2-4, 2 vvd for days 4-6, 3 vvd for days 6-8 and 4 vvd from day 8-12. Intermediate cell sampling is performed on days 1, 2, 6 and 12. Offline cell count measurements are performed on all these days, while flow cytometry for phenotyping is performed only on days 1, 6 and 12. In addition, metabolite concentrations in the perfusate are measured each day.

### Supplementary Figure 2

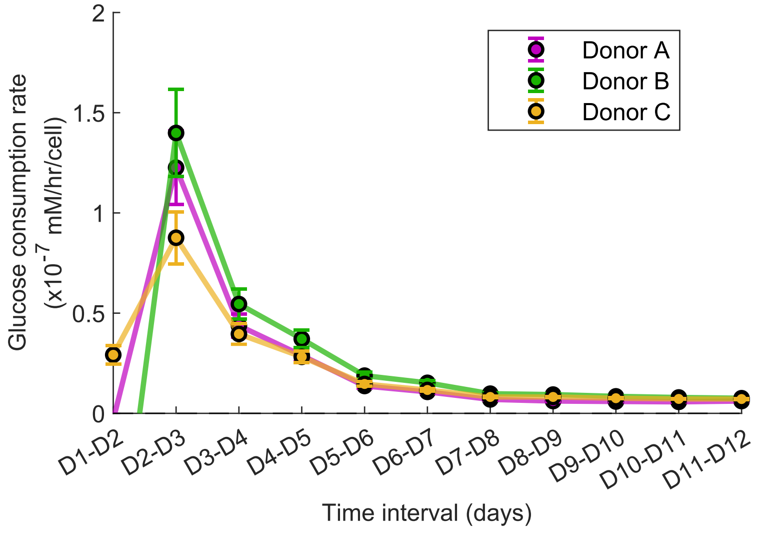


**Supplementary Figure 2. Estimated Glucose consumption rates for CAR-T cells obtained from healthy donors**. Glucose consumption rates for the three healthy CAR T cell samples. Similar to the lactate production rates in main text figure 3B, this is an average rate across a 24-hour interval, as it is based on daily offline measurements of metabolite concentrations. Error bars indicate the uncertainty arising from VCC estimation from OD values (mean ± SD). Differences can be observed between the three samples during the early days (Days 1-4) immediately following T cell activation.

### Supplementary Figure 3

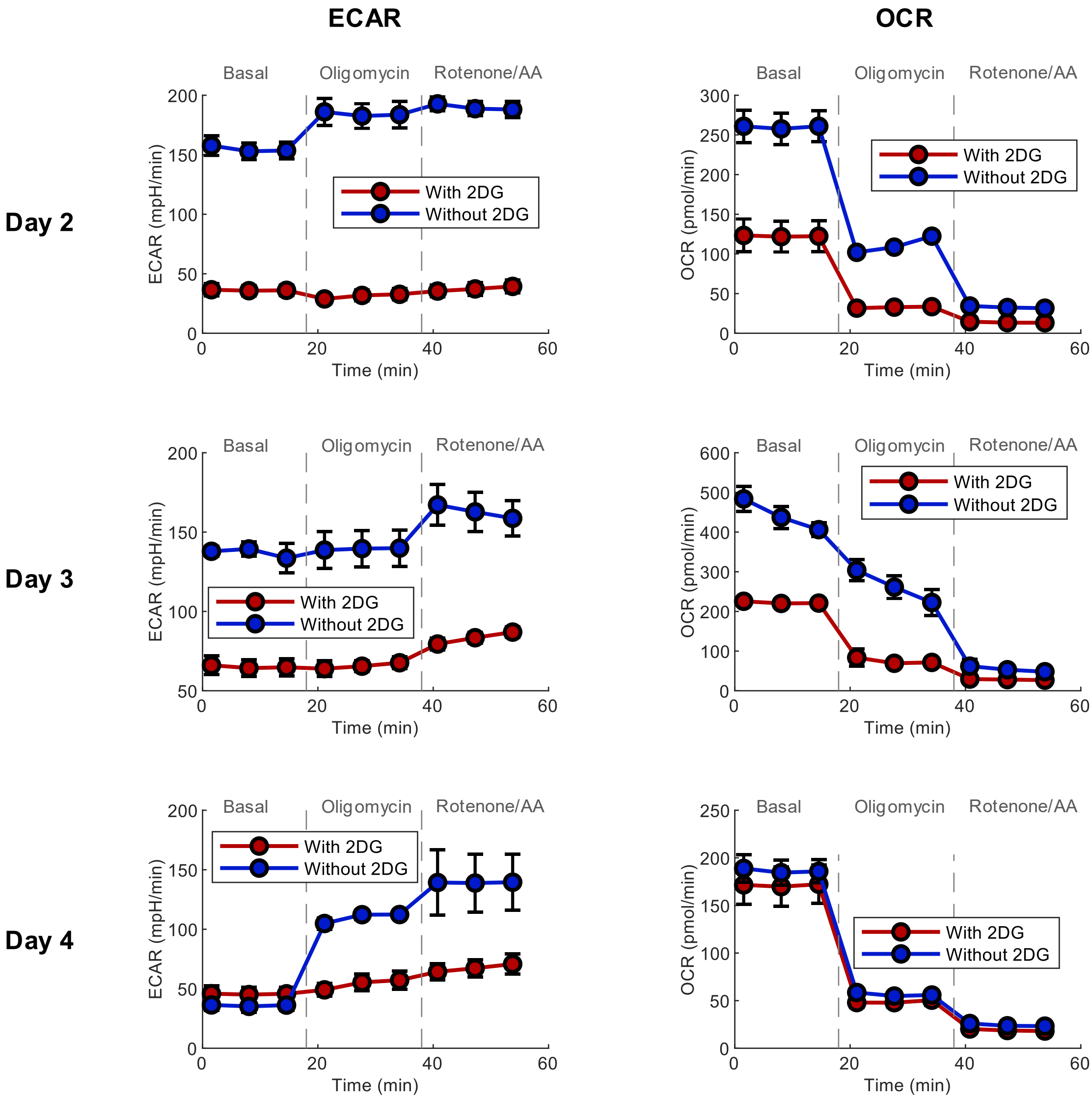


**Supplementary Figure 3.** Plots of Extracellular acidification rates (ECAR) and Oxygen consumption rates (OCR) obtained from the Seahorse extracellular flux (XF) metabolic analyzer for cell samples from Days 2, 3 and 4, in both native and 2-DG conditions (mean ± SD).

### Supplementary Figure 4

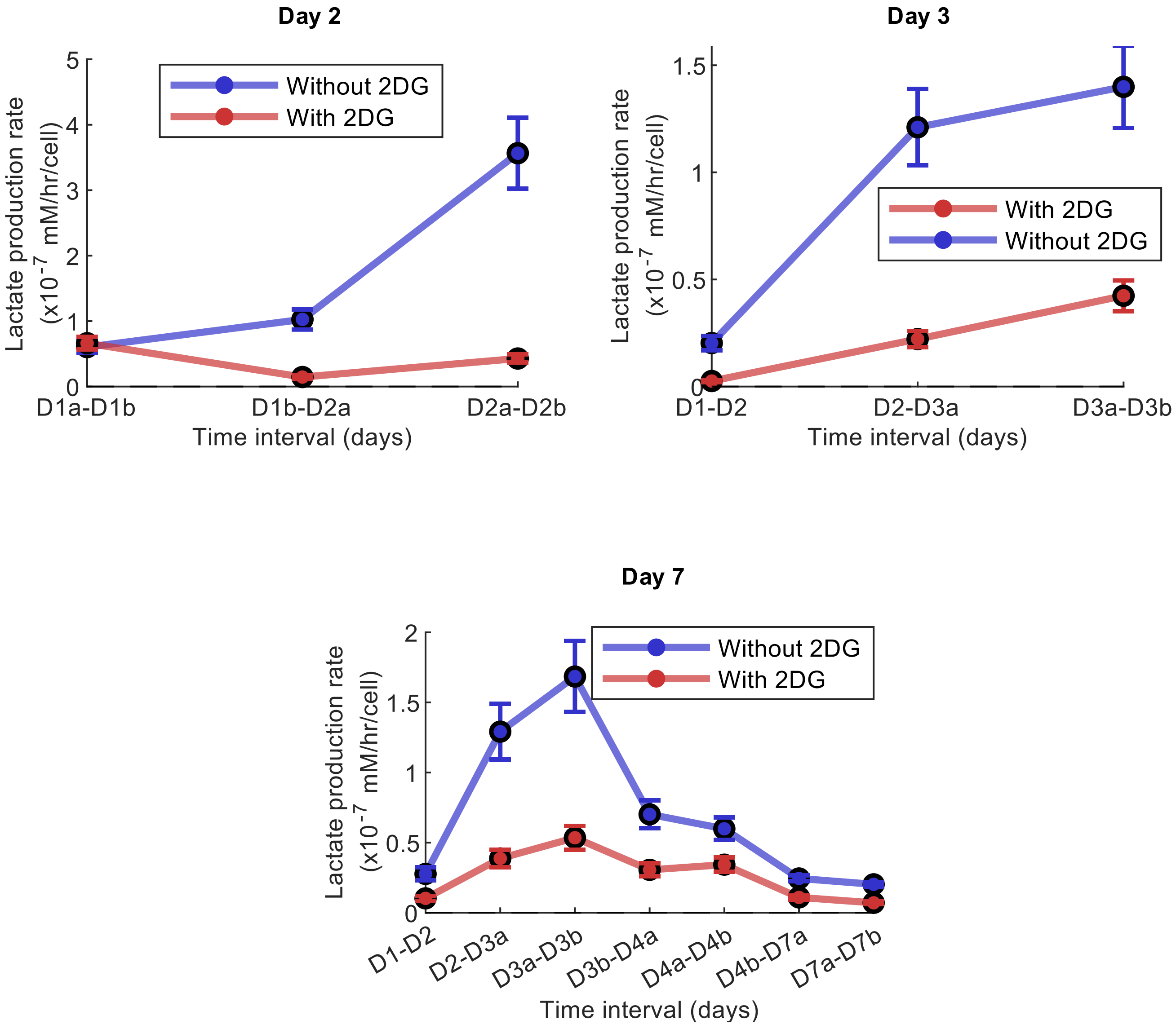


**Supplementary Figure 4. Estimated Lactate Production rates for cell samples harvested on days 2, 3 and 4.** Average lactate production rates (estimated) per cell for the interval between two consecutive offline metabolite measurements, for both the native case (blue) and the 2-DG case (red). Error bars indicate the uncertainty arising from VCC estimation from OD values (mean ± SD).

### Supplementary Figure 5

Day 2

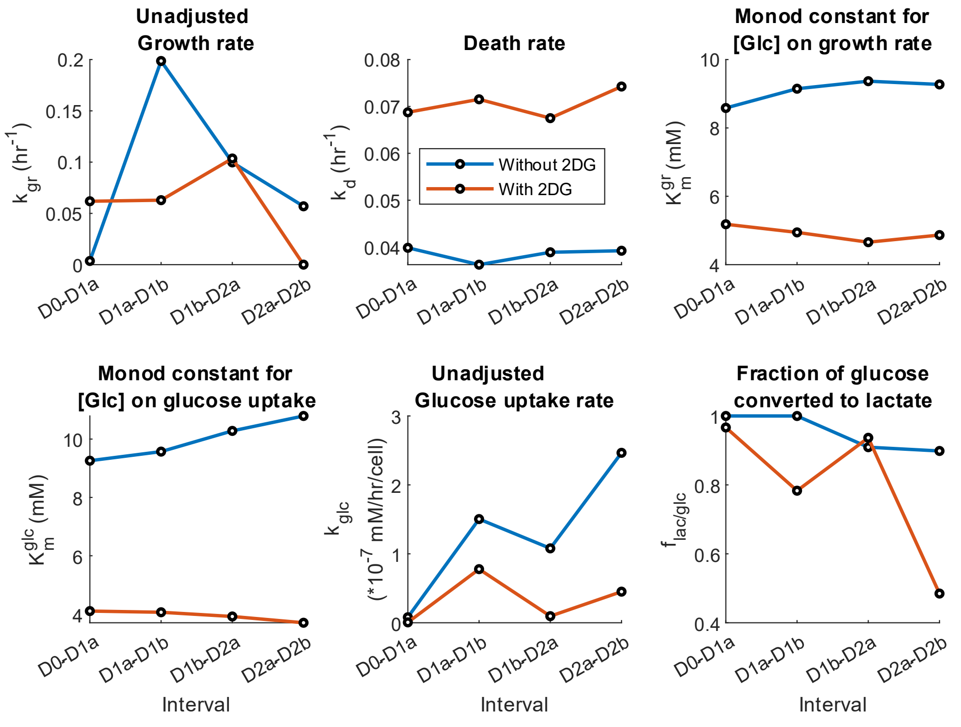


Day 3

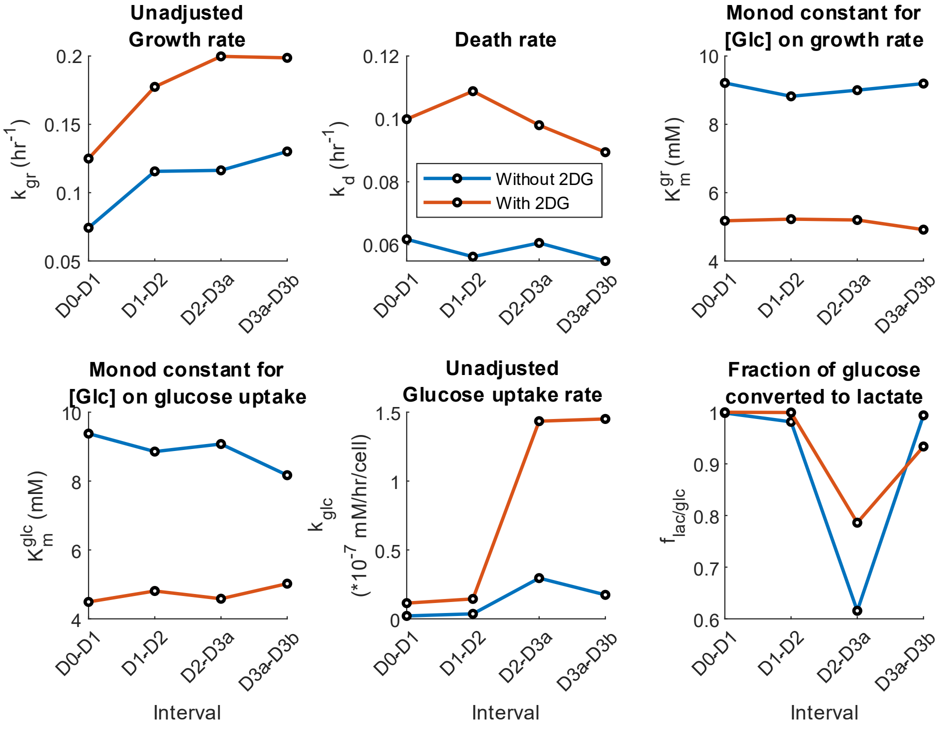

**Supplementary Figure 5. Rate parameter estimates for downsampled MBR datasets.** Best-fit estimates for the daily average of six rate parameters/variables from the full ODE model (refer to Figure 2C), for both the native (blue) and 2DG (red) conditions, for cells harvested on days 2 and 3. Corresponding figures for cells harvested on day 7 can be found in main text (Figure 5C).

### Supplementary Figure 6

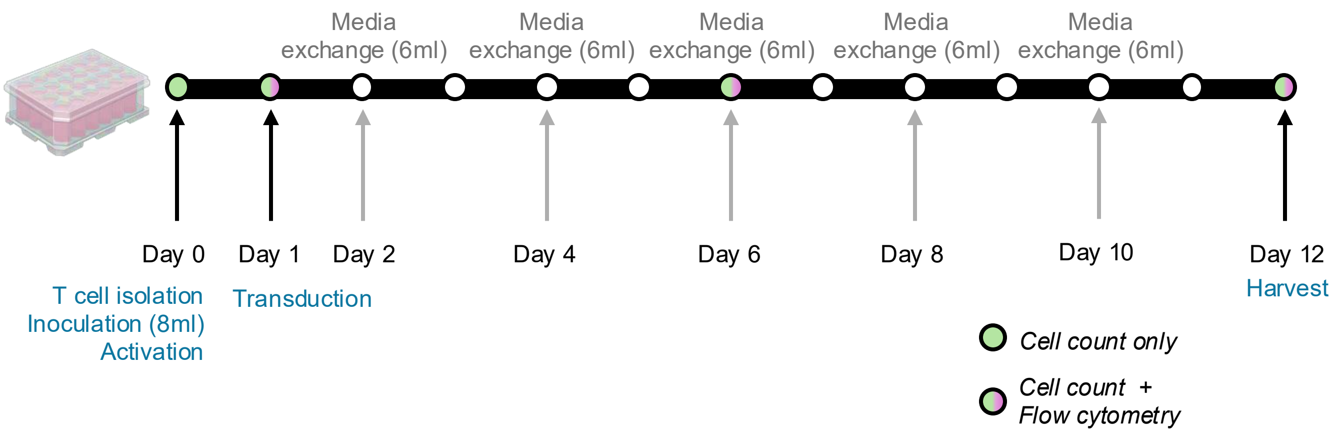


**Supplementary figure 6. Experimental design for G-Rex well plate manufacturing of CAR T cells from healthy donors.** PBMCs extracted from healthy donors are processed to obtain 2 million CD3+ T cells, that are inoculated into the G-Rex 24 well-plate (8ml media volume) on Day 0 and activated using Dynabeads. On Day 1, lentiviral transduction of CD19 CAR constructs is performed. The cells are then expanded in the well plate until Day 12 when they are harvested. Every two days, (starting from day 2), 6ml of spent media is removed from the plate and replaced with 6 ml of fresh media. Offline cell count measurements and flow cytometry for phenotyping are performed on days 1, 6 and 12. In addition, metabolite concentrations in the perfusate are measured each day. Created in BioRender. Jagannathan, S. (2025) https://BioRender.com/nz07r84

### Supplementary Figure 7

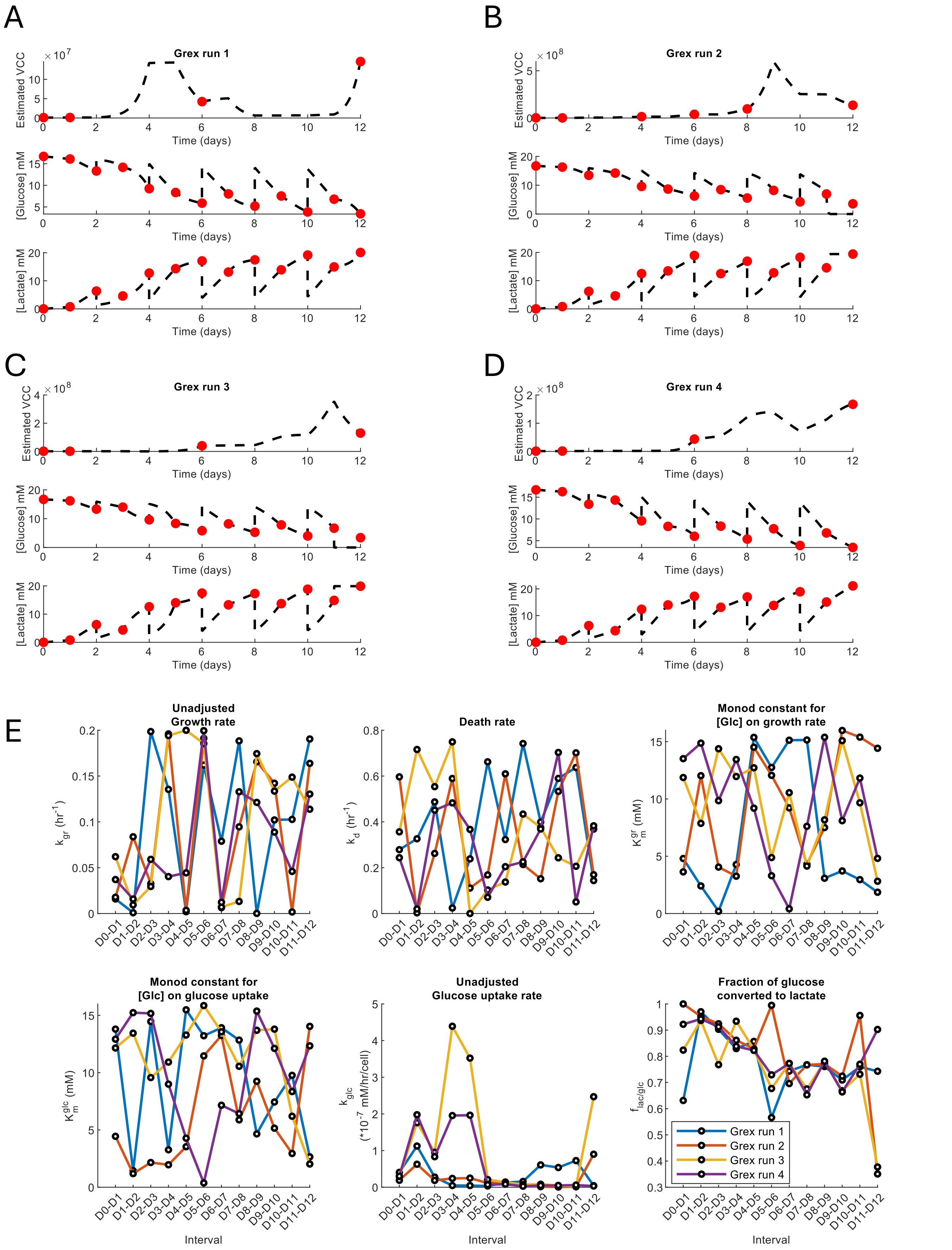

**Supplementary figure 7. Estimating rate parameters from well plate expansion of CAR T cells with sporadic cell count measurements.** (A-D) Fit between estimated and measured parameters (viable cell count, glucose and lactate concentrations) for each of the four G-Rex expansion experiments. Red points indicate measured values and dashed lines indicate computational estimation at and between the same timepoints. It can be seen that while the modelling recapitulates experimentally measured values well, the dynamics in the intervening period appear non physiological (for the estimated VCC values). This is because sporadic cell count measurements impose fewer constraints on the optimization problem during parameter estimation. This process can be improved by having more regular (or continuous) cell count measurements. (E) Best-fit estimates for the daily average of six rate parameters/variables from the full ODE model (refer to Main text Figure 2C), for each of the four G-Rex runs. The rate parameters appear to vary substantially between successive intervals, making the model less valuable for physiological interpretation in a data-poor scenario with infrequent cell count measurements

### Supplementary Figure 8

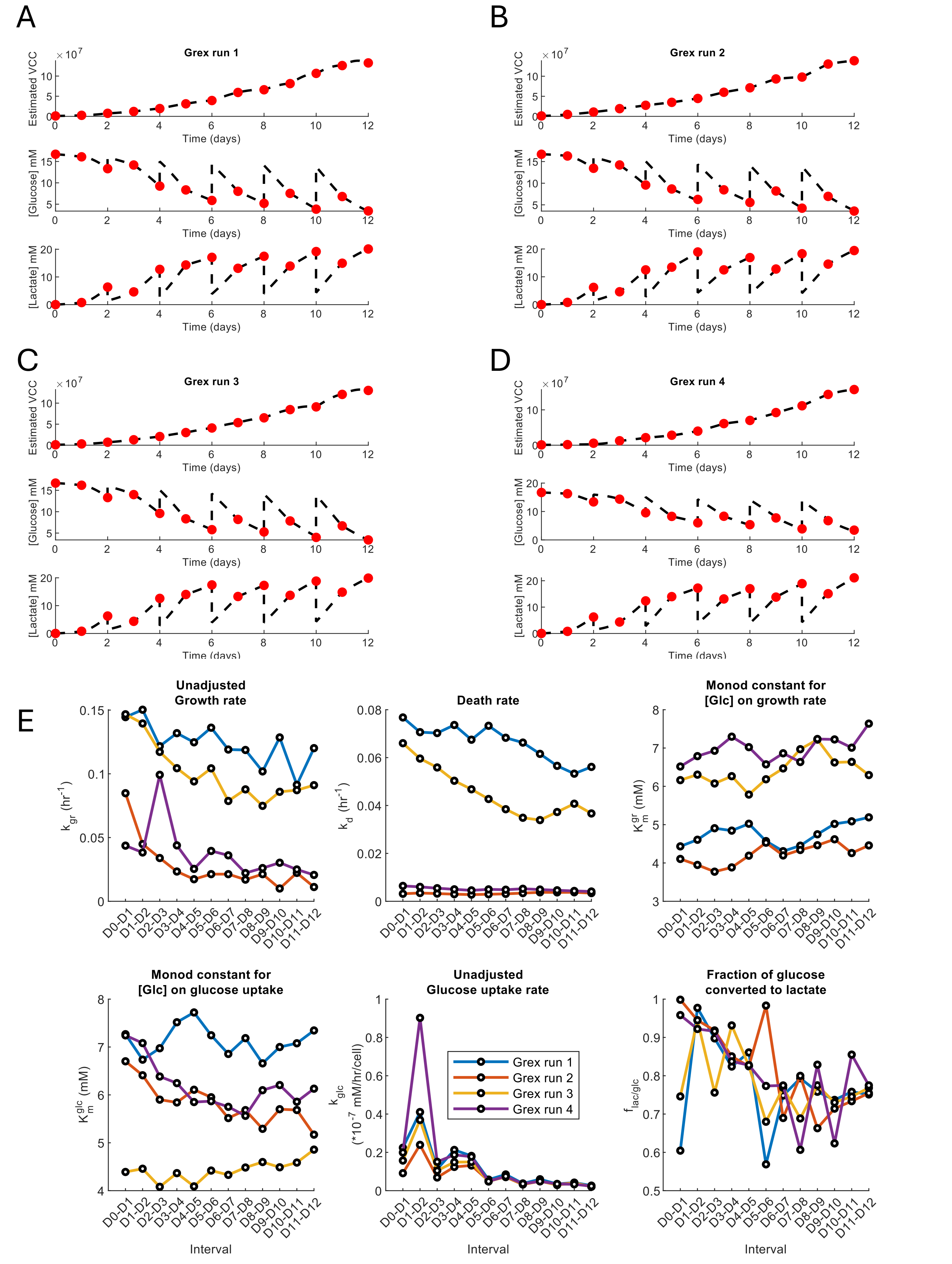

**Supplementary figure 8. Estimating rate parameters from well plate expansion of CAR T cells with daily cell count measurements.** Figure shows the same plots as Supplementary figure 7 using the same dataset, but in this case we re-estimated parameters by artificially interpolating experimental cell count values between actual measurements to simulate daily cell count measurements (instead of just days 0, 1, 6 and 12). (A-D) Fit between estimated and measured parameters (viable cell count, glucose and lactate concentrations) for each of the four G-Rex expansion experiments. Red points indicate measured (and interpolated) values and the dashed lines indicate computational estimation at and between the same timepoints. Compared to Supplementary figure 7, the dynamics of estimated cell counts and metabolites are more gradual for all runs, illustrating the improvement in modelling capabilities with regular cell sampling for batch reactor systems. (E) Best-fit estimates for the daily average of six rate parameters/variables from the full ODE model (refer to Main text Figure 2C), for each of the four G-Rex runs. While the variation in estimated rate parameters is lower than observed in Supplementary figure 7 (using the same dataset), it can be still high in some cases (fraction of glucose converted to lactate) affecting physiological interpretation.

### Supplementary Figure 9

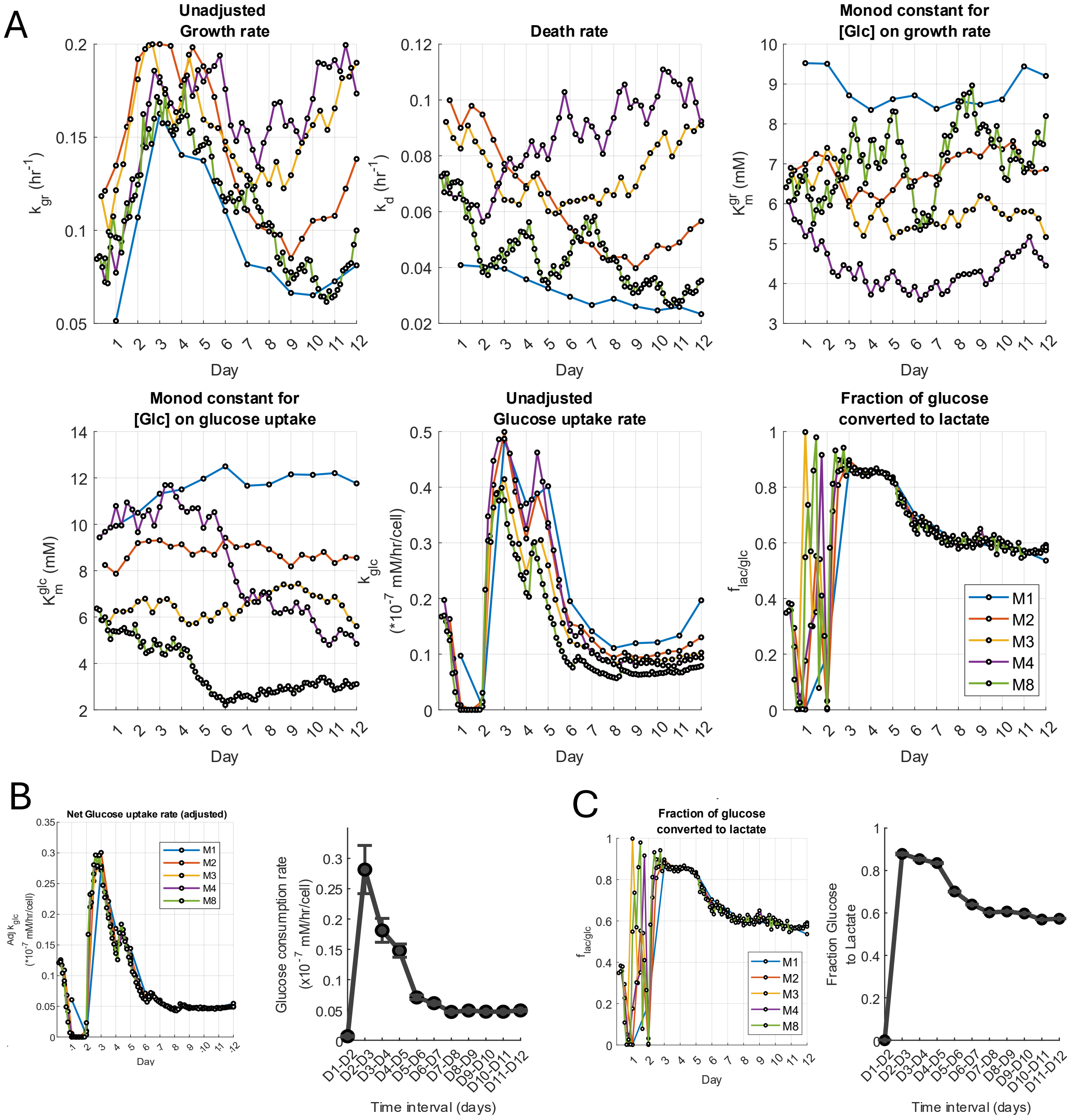


**Supplementary figure 9. Effect of greater frequency of VCC and metabolite measurements per day on estimated rate constants .** We regenerated new downsampled MBR datasets from a healthy donor run by interpolating to simulate multiple offline VCC and metabolite measurements daily (1, 2, 3, 4, or 8 measurements a day, indicated by M1, M2, M3, M4 and M8 respectively) and re-estimated growth/metabolic rates. (A) Rate parameters estimated similar to Supplementary figure 8 show wild variability in some parameters but remarkable consistency in others ($k_{glc}$, $f_{lac/glc}$), across interpolation counts. (B) Comparing adjusted $k_{glc}$ (adjusting for the effect of instantaneous glucose concentration using the Monod constant $K_m^{glc}$) results in good agreement between the estimated rates from downsampled dataset (left) and continuous OD measurements (right). (C) Similar to (B) but for fraction of glucose converted to lactate. There is very good agreement between different interpolation counts (left) and with continuous OD measurements (right) except during early timepoints

### Supplementary Figure 10

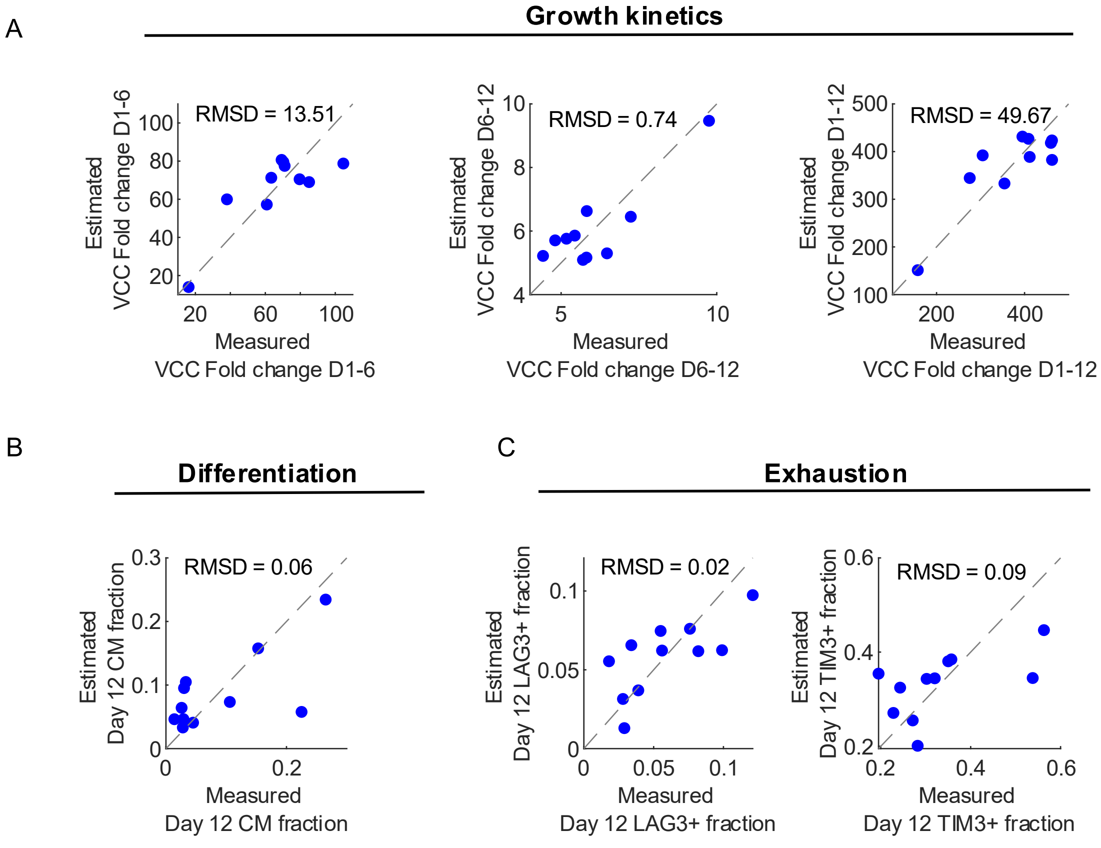


**Supplementary figure 10. Prospective prediction of final cell product attributes using early metabolic estimates.** For illustrative purposes, we performed forward prediction of final cellular attributes (growth rate, exhaustion and phenotype), using a simple linear regression model. Parameters for this regression model were estimated using the data in Main text figure 6E-G and re-applied to the same dataset to recapitulate the corresponding attribute values. Figure panels above show the concordance between the measured and predicted attribute values.

### Supplementary Figure 11

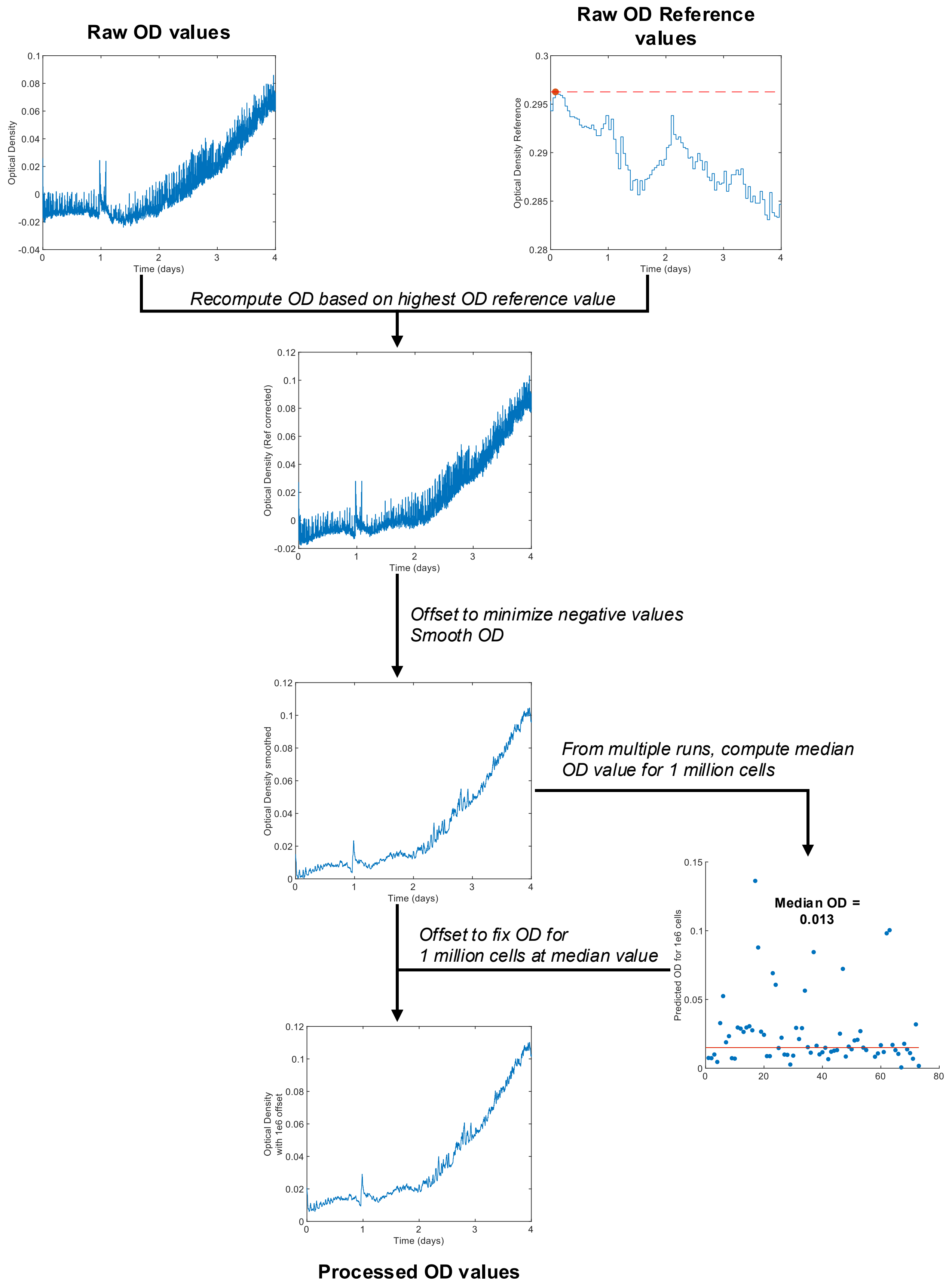

**Supplementary figure 11. Preprocessing of Optical Density values.** The optical density (OD) values from the bioreactor are pre-processed in the following manner, before ODE modelling and optimization. We first extract the raw OD values and the raw reference OD values (no cells, only media) recorded by the microbioreactor. The reference value is subject to fluctuations due to debris and other reasons. We recalculate new OD values based on the highest recorded OD reference value. An offset is then added to remove negative OD values. Finally another offset is added to adjust ODs such that the OD for 1 million cells corresponds to a fixed value. This fixed value of 0.013 is the median value for 1 million cells obtained from multiple runs of the microbioreactor. The process of OD processing is also detailed in the Methods section.

### Supplementary Figure 12

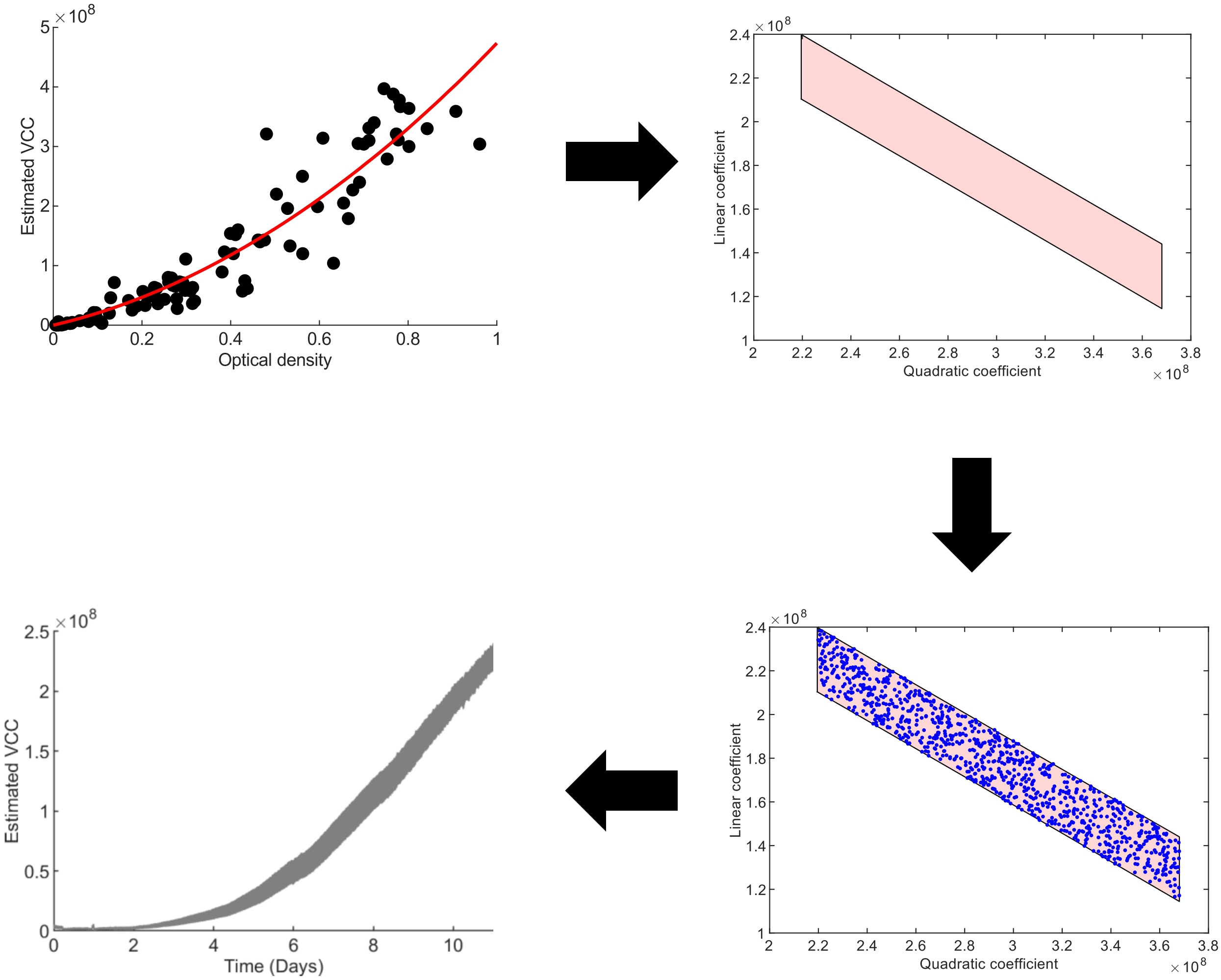


**Supplementary figure 12. Obtaining general fit between optical density and viable cell count.** We first obtain a general quadratic equation to map OD values to viable cell counts (VCCs) using paired OD and offline cell measurements from multiple bioreactor runs. The confidence intervals for the linear and quadratic coefficients of the fit are then used to obtain a feasible region in the quadratic coefficient-linear coefficient space. All combinations within this feasible region might provide equations with acceptable fits between OD and VCC. Finally we sample this feasible space to obtain 1000 equations to estimate VCCs from OD values. Using the 1000 equations gives a measure of uncertainty in the predicted viable cell counts.
